# ABP1/ABL3-TMK1 cell-surface auxin signaling directly targets PIN2-mediated auxin fluxes for root gravitropism

**DOI:** 10.1101/2022.11.30.518503

**Authors:** Lesia Rodriguez, Lukáš Fiedler, Minxia Zou, Caterina Giannini, Aline Monzer, Dmitrii Vladimirtsev, Marek Randuch, Yongfan Yu, Zuzana Gelová, Inge Verstraeten, Jakub Hajný, Meng Chen, Shutang Tan, Lukas Hoermayer, Lanxin Li, Maria Mar Marques-Bueno, Zainab Quddoos, Gergely Molnár, Tongda Xu, Ivan Kulich, Yvon Jaillais, Jiří Friml

**Author notes:** Equal contributions (Lesia Rodriguez, Lukáš Fiedler). Lead Contact: Further information and requests for resources and reagents should be directed to and will be fulfilled by the lead contact, Jiří Friml.

## Abstract

Phytohormone auxin and its directional transport mediate much of the remarkably plastic development of higher plants. Positive feedback between auxin signaling and transport is a key prerequisite for (i) self-organizing processes including vascular tissue formation and (ii) directional growth responses such as gravitropism. Here we identify a mechanism, by which auxin signaling directly targets PIN auxin transporters. Via the cell-surface ABP1-TMK1 receptor module, auxin rapidly induces phosphorylation and thus stabilization of PIN2. Following gravistimulation, initial auxin asymmetry activates autophosphorylation of the TMK1 kinase. This induces TMK1 interaction with and phosphorylation of PIN2, stabilizing PIN2 at the lower root side, thus reinforcing asymmetric auxin flow for root bending. Upstream of TMK1 in this regulation, ABP1 acts redundantly with the root-expressed ABP1-LIKE auxin receptor ABL3. Such positive feedback between cell-surface auxin signaling and PIN-mediated polar auxin transport is fundamental for robust root gravitropism and presumably also for other self-organizing developmental phenomena.

## INTRODUCTION

Plant developmental mechanisms differ fundamentally from animals. With cells encapsulated in rigid cell walls without the possibility of migration, plants mainly rely on oriented cell divisions and expansions and have the capacity to self-organize complex tissues. Being rooted in the soil, plants are also highly adapted to cope with changing environments. Much of the adaptability and self-organization is mediated by the phytohormone auxin with examples including the formation of an embryonic axis, regular arrangement of leaves and flowers, establishment of leaf vein patterns, or flexible vasculature regeneration around a wound^1^. Auxin also acts as a key endogenous signal positioning sessile plants in their environment during directional growth responses such as gravitropism and phototropism^2^. Both self-organizing development and translation of environmental signals into directional growth rely on mechanistically elusive feedback between auxin signaling and polar auxin transport^3,4^.

Directional cell-to-cell auxin transport is a plant-specific mechanism^5^ dependent upon plasma membrane-localized transporters^6,7^. Chief among these are AUX1/LAX importers^8^ and PIN auxin exporters. The latter inhabit polarized plasma membrane domains to determine vectorial auxin fluxes through tissues^9,10^. Inside cells, auxin triggers a well-studied transcriptional pathway through predominantly nuclear TIR1/AFB receptors. This leads to developmental reprogramming^11,12^ and contributes to growth regulation^13^. Historically, several auxin responses showed such rapidity that transcriptional cascades did not suffice for their explanation. While several of these were later found to also depend on TIR1/AFB receptors, other responses require extracellular (apoplastic) auxin perception^14^. This has been formalized recently as comprising AUXIN-BINDING PROTEIN1 (ABP1), ABP1-LIKEs (ABLs), and TRANSMEMBRANE KINASEs (TMKs) [ABP1/ABL-TMK] co-receptor complexes at the cell-surface^15–17^.

Sensitive phospho-proteomic pipelines recently revealed that auxin triggers a global phosphorylation response via ABP1 and TMK1^17,18^. Notably, the lack of auxin-induced phosphorylation in *abp1* and *tmk1* mutants correlates with strong defects in auxin canalization^17^, a mysterious process underlying self-organized plant development including regeneration of vasculature and formation of polarized auxin-transporting channels from a local auxin source. Canalization also requires TIR1/AFB receptors^19^, suggesting that both intracellular and apoplastic signaling contribute to auxin feedback regulation of PIN-dependent auxin transport. This is consistent with computational predictions exploring potential mechanism of PIN polarization by auxin feedback^20^, however, no such mechanism linking auxin signaling and transport has been discovered.

Feedback between auxin and its transport has also been proposed for gravitropic root bending. Contrary to canalization, this would not involve adjustments of PIN polarity but rather the stabilization of the root-specific PIN2 transporter^21^. Among the latest novelties in the quest of plants to grow upright is evolution of fast root gravitropism, which was enabled by functional innovations in the PIN2 protein^22^. Fast gravitropism occurs through directional auxin transport from the site of gravity perception towards the elongation zone where growth response takes place. After gravity sensing in the columella at the root tip^23^, auxin flux becomes redirected to the lower root side^24,25^. This initial asymmetry is then propagated by AUX1- and PIN2-mediated transport^26,27^ from the root tip to the elongation zone and translated into root bending through local inhibition of cell elongation^28^.

During the gravitropic response, PIN2 distribution itself becomes asymmetric with increased and decreased PIN2 stability at the lower and upper root sides, respectively^21,29^. Such lateral PIN2 gradient not only propagates but also reinforces the initial root tip auxin asymmetry, contributing to the robustness of root bending as well as to the fine-tuning of gravitropism by other hormonal cues^30^. How auxin regulates its own transport via PIN2 in the context of root gravitropism remains an outstanding question.

In search of a possible mechanism for cell-surface auxin signaling effect on auxin transport, we mined the auxin-inducible, ABP1-TMK1-mediated phospho-proteome and identified enrichment of PINs with PIN2 as the main target. We find that an auxin-induced interaction of TMK1 with PIN2 and its phosphorylation are directly responsible for the PIN2 gradient that reinforces gravitropic root bending. This pathway perceives auxin through the newly identified root-expressed ABL3 receptor acting redundantly with ABP1. Our findings identify a direct mechanism for feedback regulation between auxin signaling and auxin transport.

## RESULTS

### ABP1-TMK1 cell-surface auxin signaling induces phosphorylation of PIN auxin transporters

To identify components of feedback regulation of auxin transport downstream of ABP1-TMK1 auxin signaling, we took advantage of a rapid phospho-proteomic dataset (100 nM IAA, 2 min) recorded in roots of *Arabidopsis thaliana* (Arabidopsis) wild-type (WT) or the respective mutants^17^. We queried proteins concurrently hypo-phosphorylated in both *abp1-TD1* and *tmk1-1* mutants for molecular function using a gene ontology (GO) analysis. When partitioned by significance, the most dominant terms were rather general and included “binding” or “protein binding”. On the other hand, partitioning significant terms by effect size (fold enrichment) always recovered “auxin efflux transmembrane transporter activity” as the most strongly enriched GO term (Figure S1A and S1B). Inspection of the corresponding enriched phospho-proteins showed the presence of PINs and ABCB/ABCG transporters. We further focused on PINs as dominant auxin transporters with many established developmental roles.

There were in total nine PIN phospho-peptides (phospho-sites) significantly downregulated in both *abp1-TD1* and *tmk1-1* (Figure 1A and Figure S1C). To verify the genetic specificity of these results, we examined in parallel a recent matched auxin phosphorylation dataset^18^ from the mutant of the intracellular AFB1 auxin receptor^31^. Except for PIN1^S337^, none of the ABP1-TMK1-dependent PIN phospho-sites were deregulated in *afb1-3* (Figure 1A). This suggests that auxin activates PIN phosphorylation specifically through the cell-surface ABP1-TMK1 module independently of intracellular TIR1/AFB signaling. Two of the nine PIN phospho-sites mapped to PIN1, five to PIN2, and two to PIN3 (Figure 1A). Notably, all these targeted hydrophilic PIN loops, the expected location for post-translational modifications regulating PIN function^32^. The two PIN1 sites, PIN1^S271^ and PIN1^S337^, were previously ascribed to shoot functions as targets of the D6PK protein kinase^33^ and the MKK7-MPK6 module^34^, respectively. Another previously studied phospho-site was PIN2^S439^, which participates in root adaptation to varying nitrogen sources^35,36^.

**Figure 1.**
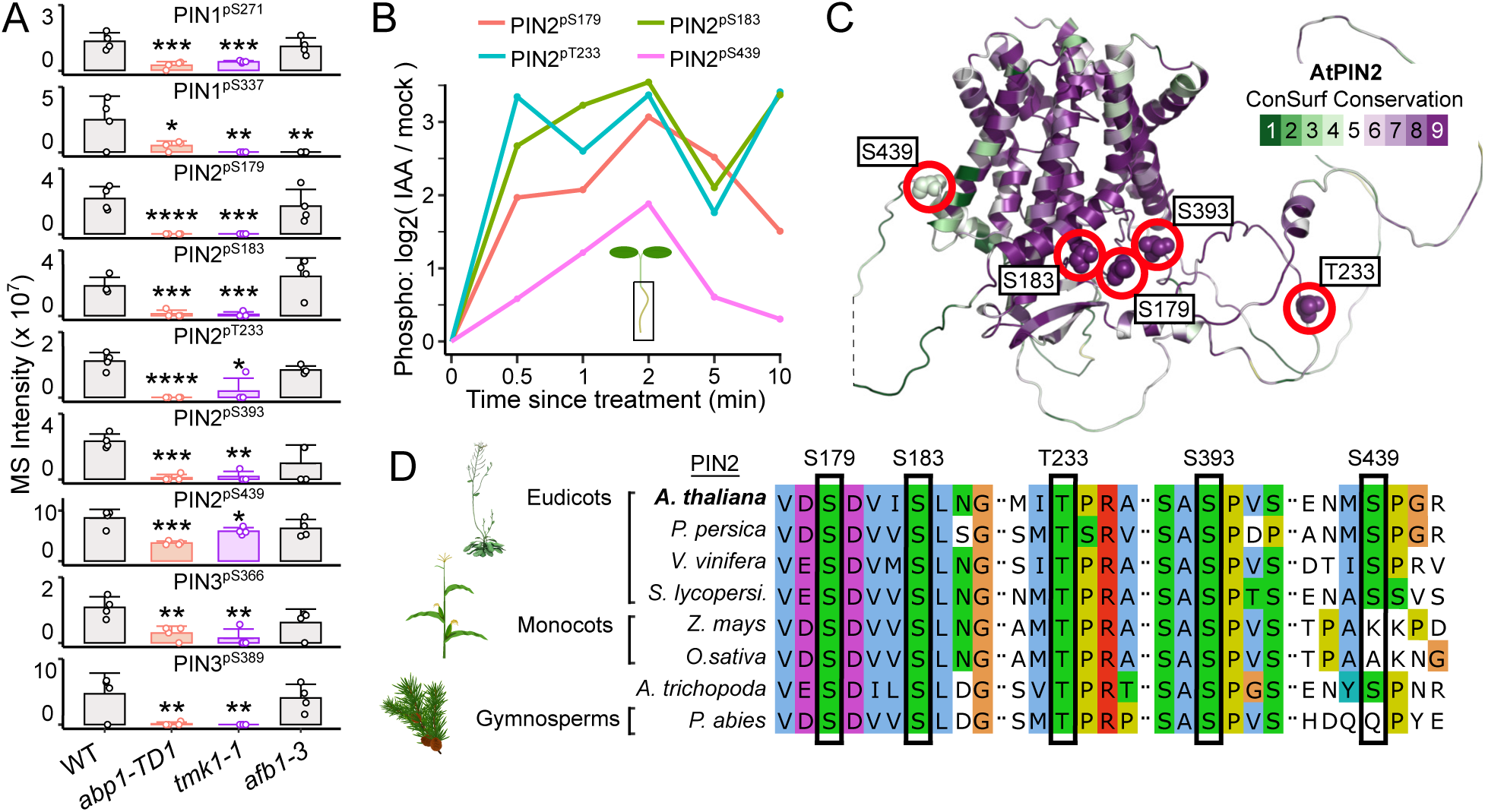
Rapid auxin phospho-response targets PINs in Arabidopsis. (A) Overview of PIN phospho-sites downregulated (FDR < 0.05) in *abp1-TD1* and *tmk1-1* auxin-treated (100 nM IAA, 2 min) roots. 4 biological replicates, Mean + SD, permutation-based t-tests with FDR-controlled p-values. *p < 0.05, **p < 0.01, ***p < 0.001, ****p < 0.0001. (B) Significant PIN2 phospho-site auxin profiles (FDR ≤ 0.01, 100 nM IAA). (C) Localization of ABP1-TMK1-dependent phospho-sites on a ConSurf conservation-colored AlphaFold2 structure of PIN2. (D) Multiple sequence alignment of eight PIN2 orthologs with a highlight of Arabidopsis ABP1-TMK1-dependent PIN2 phospho-sites.

Given that cell-surface auxin signaling mutants show perturbed phospho-proteomes even under mock conditions^17,18^, we next assessed auxin inducibility of these phospho-sites in WT roots. Notably, four out of five PIN2 phospho-sites strongly responded to 100 nM IAA within 30 seconds of treatment (Figure 1B). The PIN3^S389^ phospho-site showed a similar behavior. Conversely, while the auxin profile of PIN1^S337^ also significantly deviated from mock conditions, the site only underwent a delayed negative fluctuation (Figure S1D). This correlates with PIN1^S337^ not being specifically targeted by ABP1-TMK1 (Figure 1A). We also confirmed average to low evolutionary conservation of PIN1^S337^ (Figure S1E), altogether suggesting minor biological relevance of this particular site for the ABP1-TMK1 auxin phospho-response. When extending evolutionary conservation analysis to the remaining PIN phospho-sites, we observed moderate conservation of PIN1^S271^ and poor conservation of the auxin-inducible PIN3^S389^ (Figure S1E and S1F). On the other hand, four out of five PIN2 phospho-sites showed high ConSurf scores and perfect conservation in PIN2 orthologs from Arabidopsis to Gymnosperms (Figure 1C and 1D).

Given that rapid auxin phospho-response represents an ancient auxin pathway^18^, we next asked whether PIN phosphorylation is conserved across the green lineage. Unlike in Arabidopsis, we found no significantly regulated PIN phospho-sites in the auxin phospho-proteomes (100 nM IAA, 2 min) of two bryophytes (*Physcomitrium*, *Marchantia*) and two streptophyte algae (*Penium*, *Klebsormidium*). Conversely, the fern *Ceratopteris* showed auxin-regulated PIN phosphorylation under the same conditions^37^. This suggests a co-option of an ancestral auxin response for PIN phosphorylation after the divergence of Bryophyta from the green lineage, probably in the common ancestor of vascular plants.

These analyses identified PIN auxin transporters, particularly PIN2, as prominent targets of ultrafast ABP1-TMK1-mediated auxin phospho-response, representing a recent evolutionary innovation.

### ABP1-TMK1-dependent PIN2 phospho-sites are crucial for PIN2 stability and root gravitropism

In further investigations, we focused on PIN2, as it was most extensively targeted, its phosphorylation strongly responded to auxin, and it showed remarkable conservation at the majority of its phospho-sites.

To test the physiological relevance of ABP1-TMK1-dependent PIN2 phosphorylation, we mutated the five candidate phospho-sites (Figure 2A and 2B) to either aspartate or alanine and introduced the resultant phospho-variants in the agravitropic *eir1-4* mutant^21^ under the native PIN2 promoter. This yielded *PIN2::PIN2^WT^-GFP;eir1-4* (*PIN2^WT^-GFP*), *PIN2::PIN2^5-MIMIC^-GFP;eir1-4* (*PIN2^5-MIMIC^-GFP*) and *PIN2::PIN2^5-DEAD^-GFP;eir1-4* (*PIN2^5-DEAD^-GFP*). Given the rapidity of the auxin effect on PIN2 phosphorylation (Figure 1B), we specifically focused on early stages of gravitropic root bending. While *PIN2^WT^-GFP* complemented *eir1-4* close to WT levels, the phospho-mimic *PIN2^5-MIMIC^-GFP* provided only partial rescue (Figure 2C); an effect highly reproducible among independent lines. The phospho-dead *PIN2^5-DEAD^-GFP* showed a weaker effect, which was pronounced during the first two hours of bending and then slowly dissipated (Figure 2C). Interestingly, the *PIN2^5-MIMIC^-GFP* phenotype extended beyond early gravitropic stages and was apparent even 12 hours after gravistimulation (Figure S2A), suggesting that chronic ABP1-TMK1-like phosphorylation of PIN2 strongly perturbs root gravitropism. These results collectively demonstrate the importance of ABP1-TMK1-dependent phospho-sites for the physiological function of PIN2 in root gravitropism.

**Figure 2.**
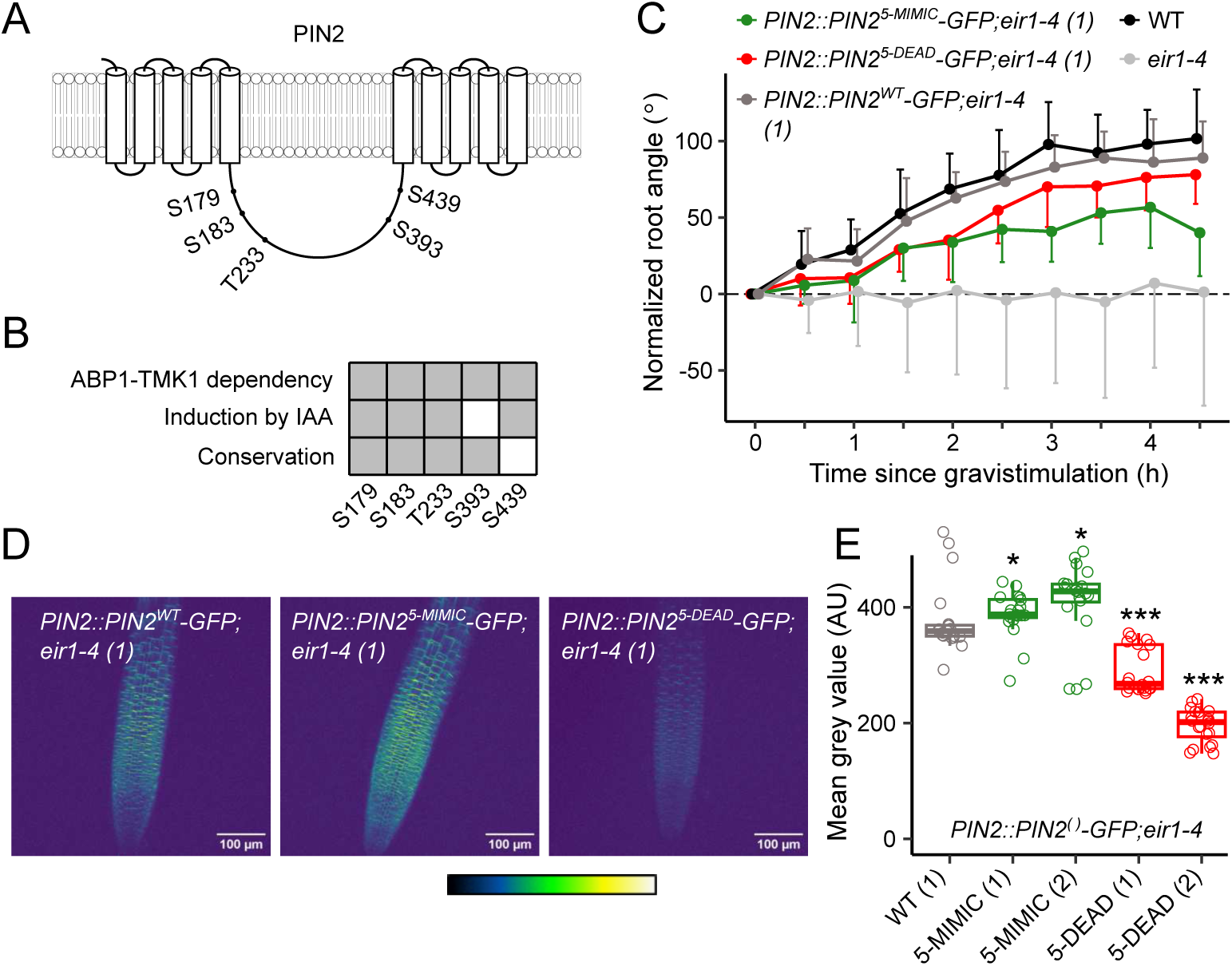
ABP1-TMK1-dependent PIN2 phospho-sites for gravitropism and PIN2 stability. (A) Schematic of ABP1-TMK1-dependent phospho-sites mapped on the PIN2 hydrophilic loop. (B) Schematic summarizing properties of the studied PIN2 phospho-sites. (C) Root gravitropism of PIN2-GFP phospho-lines on medium with sucrose (1 %). Mean + or – SD. (D) Representative maximum intensity projection images of PIN2-GFP phospho-lines. (E) Quantification of GFP signal from (D). Kruskal-Wallis analysis followed by Holm-corrected Wilcoxon rank sum tests relative to WT. *p < 0.05, **p < 0.01, ***p < 0.001, ****p < 0.0001.

Phosphorylation of PIN2 by AGC3 kinases at PIN2^S237^, PIN2^S258^, and PIN2^S310^ was previously established as regulating polar PIN2 localization^38^. However, the five ABP1-TMK1-dependent PIN2 phospho-sites were non-overlapping (Figure 2A). Indeed, we observed stereotypical polarity of apical PIN2 in epidermal cells and basal PIN2 in young cortical cells in our *PIN2^5-MIMIC^-GFP* and *PIN2^5-DEAD^-GFP* phospho-lines (Figure S2B). While the PIN2 phospho-lines showed no obvious polarity defects, we did observe reproducible differences in their GFP signal intensity. These were stronger than insertion-dependent variation when selecting T1 transformants and were also apparent in independent, single-insert, GFP-positive phospho-lines (Figure 2D). The *PIN2^5-MIMIC^-GFP* variants showed occasional weak stabilization compared to *PIN2^WT^-GFP* (Figure 2D and 2E). On the other hand, *PIN2^5-DEAD^-GFP* lines showed a highly consistent strong destabilization compared to *PIN2^WT^-GFP* roots (Figure 2D and 2E). Isolation of lines with comparable transgene mRNA levels unequivocally ruled out insertion-dependent expression artifacts and established the loss-of-phosphorylation stability defect as post-transcriptional (Figure S2C and S2D).

Altogether, our results show the relevance of ABP1-TMK1-dependent PIN2 phospho-sites for steady-state PIN2 stability and root gravitropism, suggesting a role of cell-surface auxin signaling in these processes.

### Root-expressed ABL3 auxin receptor acts redundantly with ABP1 in root gravitropism

Next, we investigated the genetic basis of PIN2 phosphorylation by cell-surface auxin signaling. It recently became recognized that apoplastic auxin perception shows multi-level redundancy^39,40^. This includes a presumably abundant pool of poorly understood ABP1/ABL auxin receptors communicating with four possible TMKs, together activating global phosphorylation reprogramming of the cellular proteome^15,17,18^. Although we identified PINs as major phospho-targets of this signaling pathway (Figure 1A), the precise composition and redundancy of the upstream auxin signaling complexes remain elusive.

TMKs form a redundant family with single mutants having rather subtle phenotypes and higher-order mutants showing strong defects in growth and development^41^. To study TMK expression in roots we used global transcriptomic data and generated *TMK1::GUS*, *TMK2::GUS*, *TMK3::GUS,* and *TMK4::GUS* lines reporting the corresponding promoter activities. The dominant family member highly expressed in roots was *TMK1*, followed by *TMK3* and *TMK4* with lower expression levels (Figure S3A-D). To examine the role of TMK1 in root gravitropism, we performed sensitive phenotyping of the *tmk1-1* mutant (Figure 3A, Figure S3F, and Figure S4D). This revealed an early root-bending defect that was complemented by a *TMK1::gTMK1-GFP* construct (Figure S3F). These data support TMK1 as the dominant TMK upstream of PIN2 phosphorylation.

**Figure 3.**
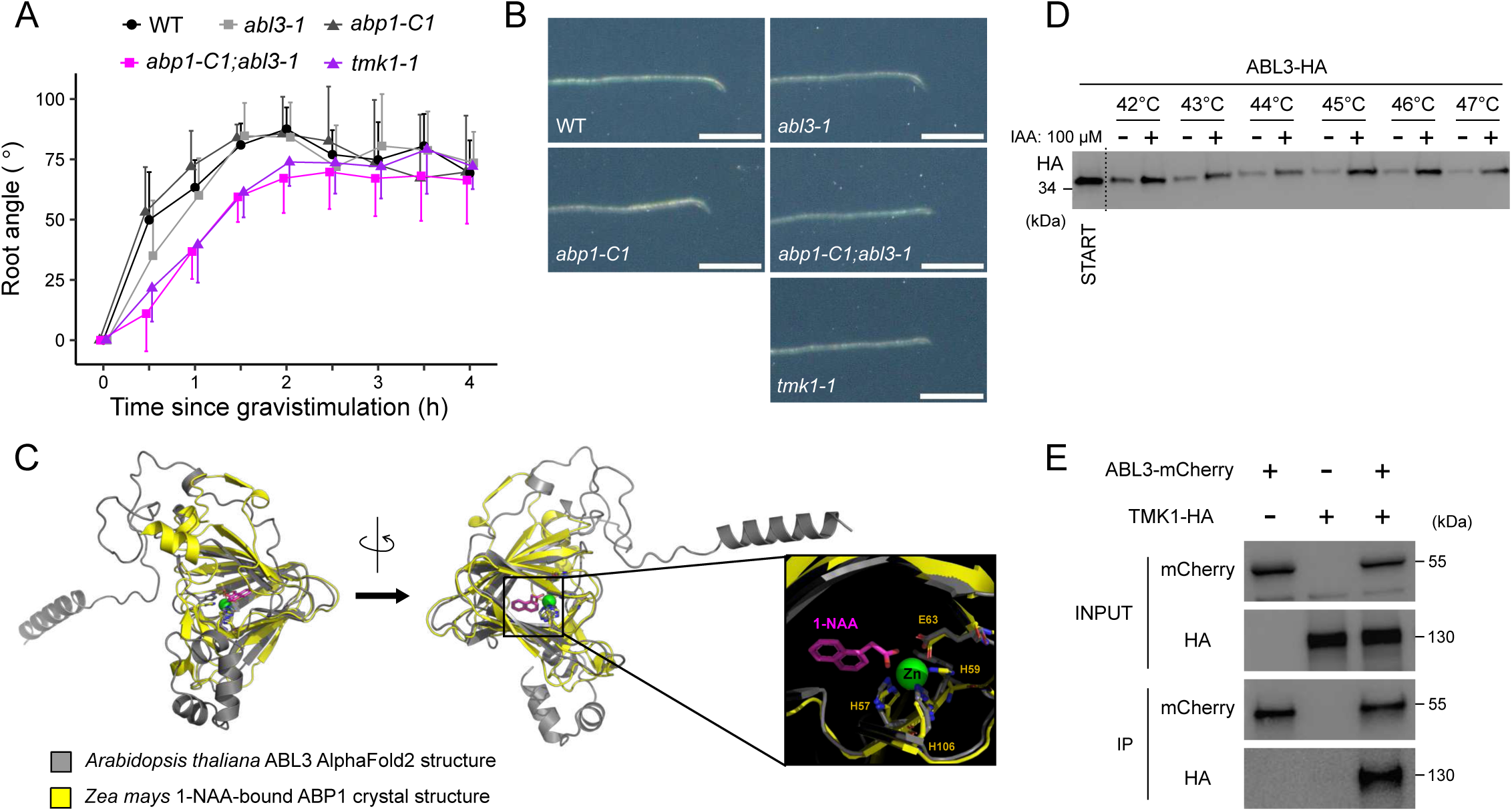
ABP1/ABL3-TMK1 signaling during early root gravitropism. (A) Mutant root gravitropism profiles on medium without sucrose. Mean + or – SD. (B) Representative images of mutant roots gravistimulated for 1 hour. The full set of images is shown in Fig. S4E. Scale bar, 100 µm. (C) Superimposition of Arabidopsis ABL3 AlphaFold2 structure with 1-NAA-bound maize ABP1 crystal structure highlighting a potential auxin-binding cavity of Arabidopsis ABL3. (D) CETSA assay on *35S::ABL3-HA* Arabidopsis root protoplasts in the presence of 100 µM IAA. (E) Co-immunoprecipitation from tobacco leaves of TMK1-HA with ABL3-mCherry but not with anti-mCherry beads alone.

Unlike *tmk1-1*, the well-established *abp1* mutant lines (*abp1-C1*, *abp1-TD1*) do not show any appreciable defects in gravitropism^42^. Nevertheless, complementation of *abp1-TD1* by native expression of an auxin-binding-deficient ABP1 variant exerts a dominant negative effect on root gravitropism, indicating the existence of unknown redundant ABLs interacting with TMK1 in the root^15^. A recent report^15^ described the redundant action of ABP1 with two auxin receptors, ABL1 and ABL2. The *abp1;abl1;abl2* triple mutant shows normal gravitropism, however, consistent with the predominantly shoot-specific expression of *ABL1* and *ABL2* (Figure S4A).

Sensitive gravitropic phenotyping led us to identify a T-DNA insertion knock-out of an ABL1/ABL2 paralog, which we named ABP1-LIKE 3 (ABL3). The *abl3-1* mutation phenocopied the early root gravitropism defects of *tmk1-1* but only in a double mutant constellation with either *abp1-C1* (Figure 3A and 3B) or *abp1-TD1* (Figure S4D and S4E). The double mutant phenotype was reproduced with an independent T-DNA insertion line, *abl3-2* (Figure S4F). We confirmed *ABL3* (AT4G14630) expression in the root by mining public RNAseq data and using an *ABL3::GUS* line (Figure S4A-C). The ABL3 protein encompasses 222 residues and does not harbor a KDEL endoplasmic reticulum retention sequence. Superimposition of Arabidopsis ABL3 AlphaFold2 structure with the 1-NAA-bound maize ABP1 crystal structure revealed a potential auxin-binding cleft in AtABL3 (Figure 3C). This also highlighted that ABL3 conforms to the ancient cupin fold of ABP1^43^. Importantly, ABL3 showed perfect conservation of three metal-coordinating residues known to be indispensable for auxin binding in ABP1, ABL1, and ABL2 (Figure 3C). The sequence surrounding these residues resembled ABL1 and ABL2 more than ABP1 (Figure S4G), as expected from members of the same GLP family^16^.

Next, we tested whether ABL3 binds auxin using a cellular thermal shift assay (CETSA) followed by western blotting. The natural auxin IAA conferred protection from thermal denaturation on ABL3-HA in protein extracts from Arabidopsis root protoplasts transformed with *35S::ABL3-HA* (Figure 3D). Likewise, IAA protected ABL3-6xHIS-3xFLAG (or ABL3-HF) in protein extracts from Arabidopsis seedlings stably transformed with *35S::ABL3-HF* (Figure S4H). These results qualify ABL3 as an auxin-binding protein.

To transmit signals from auxin-bound ABL3, TMK1 would be expected as an ABL3 interaction partner. Indeed, in tobacco leaves, TMK1-HA co-immunoprecipitated with ABL3-mCherry but not with anti-mCherry beads alone (Figure 3E). Reciprocally, we further confirmed this interaction in Arabidopsis root protoplasts where ABL3-HA co-immunoprecipitated with TMK1-mCherry but not with anti-mCherry beads alone (Fig. S4I).

Thus, we identified ABL3 as a root-expressed auxin-binding protein interacting with TMK1 and acting redundantly with ABP1 in root gravitropism. These observations are consistent with the notion that the cell-surface ABP1/ABL3-TMK1 module represents a root-specific pathway targeting PIN2 phosphorylation for early stages of gravitropic root bending.

### Exogenous and endogenous auxin activates TMK1 and downstream ROP signaling in roots

Despite recent progress, the cellular and molecular readouts of cell-surface TMK1-dependent auxin signaling remain poorly established. Previous data showed that the cytoplasmic part of TMK1 harbors an ABP1-dependent phospho-site^17^. Furthermore, the TMK1 kinase domain shows a capacity to auto-phosphorylate^44^, and research on other leucine-rich repeat receptor-like kinases (LRR-RLKs) suggests that phosphorylation of their cytoplasmic domains leads to LRR-RLK activation^45,46^.

Therefore, we examined TMK1 phosphorylation in response to auxin. We immunoprecipitated TMK1-FLAG from auxin-treated (IAA, 10 nM, 1 hour) *TMK1::TMK1-FLAG*;*tmk1-1* roots. After confirming successful IP with an anti-FLAG antibody, we stripped and re-probed the membranes with a Phos-tag Biotin Probe that coordinates tetrahedral phosphate moieties. We observed significant induction of TMK1 phosphorylation by auxin, presumably corresponding to increased TMK1 activity (Figure 4A and S5A).

**Figure 4.**
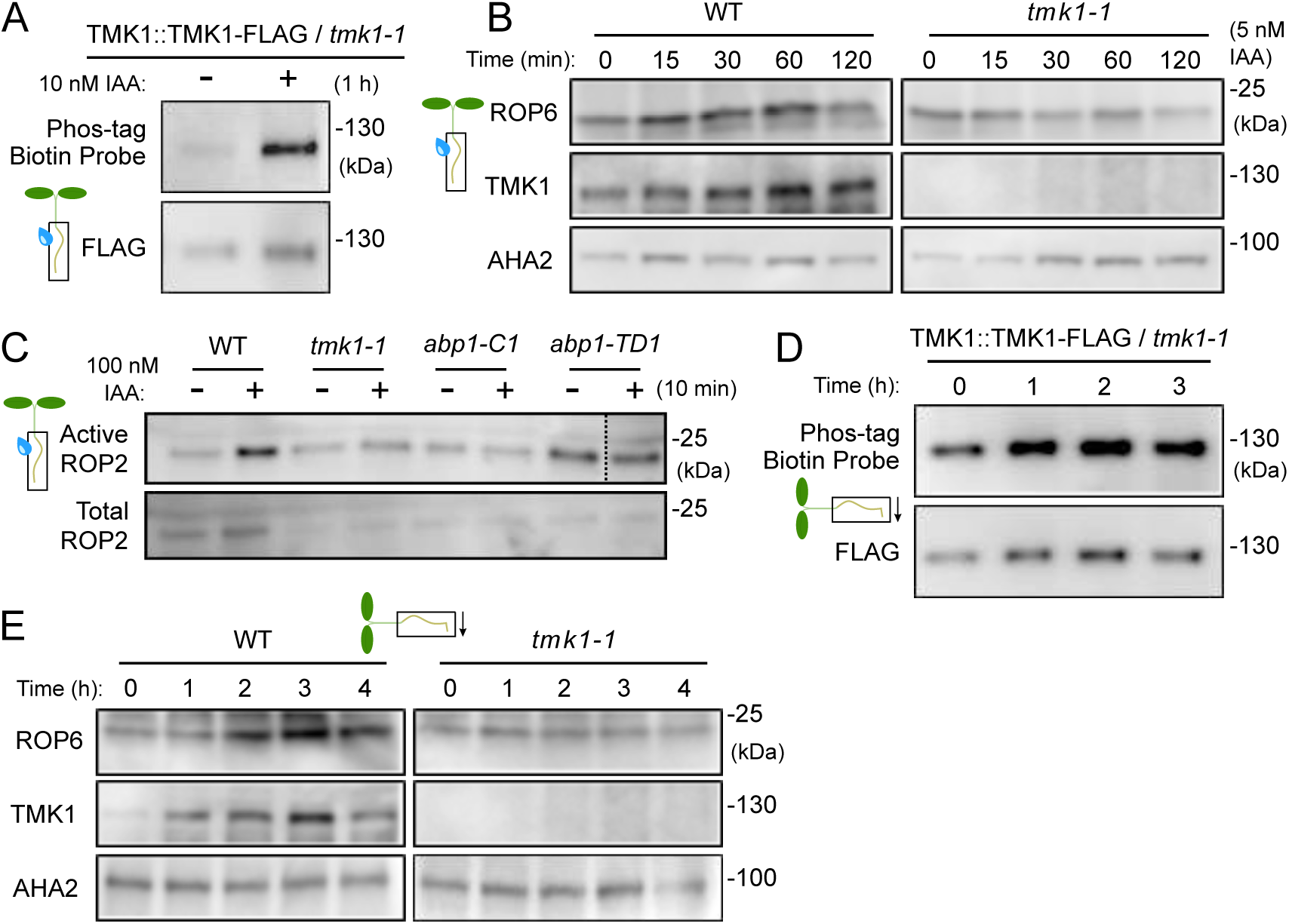
Auxin-induced activation of TMK1 and downstream ROP signaling in roots. (A) Auxin effect on TMK1 phosphorylation in *TMK1::TMK1-FLAG;tmk1-1* roots assayed through TMK1-FLAG immunoprecipitation and Phos-tag Biotin Probe analysis. Refer to Fig. S5A for quantification of three experimental replicates. (B) Auxin effect on ROP6 and TMK1 levels in the WT or *tmk1-1* root microsomal protein fractions. Refer to Fig. S5B for quantification of three experimental replicates. (C) Auxin effect on ROP2 activation assayed by native ROP pulldown from roots of the indicated genotypes. Empty well was edited out from the upper blot for visualization purposes. Refer to Fig. S5D for unedited blot image. (D) Gravistimulation effect on TMK1 phosphorylation in *TMK1::TMK1-FLAG;tmk1-1* roots assayed through TMK1-FLAG immunoprecipitation and Phos-tag Biotin Probe analysis. Refer to Fig. S5E for quantification of three experimental replicates. (E) Gravistimulation effect on ROP6 and TMK1 levels in the WT or *tmk1-1* root microsomal protein fractions. Refer to Fig. S5F for quantification of three experimental replicates.

As a downstream response, we investigated the root-specific activation of small GTPases from the ROP family implicated downstream of ABP1/ABL-TMKs. Previous ROP activation assays relied extensively on ROP overexpression, used the synthetic auxin 1-NAA, and were usually performed with leaf tissue^15,47,48^. We specifically asked if the natural auxin IAA activates ROPs in roots under non-overexpressing conditions. Immunoblotting microsomal protein extracts from auxin-treated (IAA, 5 nM, 0–120 minutes) roots with an anti-ROP6 antibody revealed auxin-induced enrichment of ROP6 in WT but not *tmk1-1* (Figure 4B and S5B). Given that membrane association is a prerequisite for ROP activation^49^, enrichment in the microsomal fraction likely reports TMK1-dependent ROP6 activation by auxin. Interestingly, we also observed that auxin stabilized TMK1 itself (Figure 4A-B and S5B) but did not induce *TMK1* mRNA over time (IAA, 10 or 100 nM, 0–120 minutes: Figure S5C). Such TMK1 stabilization at the membrane might be related to the recently reported auxin-mediated TMK1 nano-clustering effect^50^.

We next used an orthogonal method to study ROP activation by auxin. As usual for small GTPases, only GTP-bound (active) but not GDP-bound (inactive) ROP proteins engage in protein-protein interactions with their effectors. We purified the Cdc42/Rac-interactive binding motif (CRIB) domain of the RIC1 ROP effector from bacteria and used it to pull down active ROPs from native root protein extracts. Immunoblotting with an anti-ROP2 antibody revealed a strong auxin-induced (IAA, 100 nM, 10 minutes) ROP2 activation in WT but much weaker activation in *tmk1-1*, *abp1-C1*, or *abp1-TD1* roots (Figure 4C and S5D). This confirms that auxin activates root ROP2 through the ABP1-TMK1 module.

To test whether the above observations remain valid also when auxin levels are changed endogenously, we repeated TMK1-FLAG immunoprecipitation followed by a Phos-tag Biotin Probe blotting on gravistimulated roots. This revealed a significant increase in TMK1 phosphorylation (Figure 4D and S5E), indicating that gravistimulation activates TMK1. Accordingly, gravistimulation also induced a TMK1-dependent enrichment of ROP6 in the microsomal fraction (Figure 4E and S5F).

Overall, these data show that both exogenous and endogenous manipulations of auxin levels promote TMK1 phosphorylation and activation of downstream ROP GTPase signaling.

### Asymmetric activation of TMK1 and downstream ROP signaling in root gravitropism

Having established auxin-induced TMK1 activation in the root (see Figure 4 and S5) and the importance of ABP1-TMK1-dependent PIN2 phosphorylation for its stability and in early gravitropic root bending (see Figure 1, 2, S1, and S2), we assessed the role of these regulations in root gravitropism.

Our experiments with native ROP activation suggested auxin-responsive ROP signaling as a suitable proxy for TMK1 activity in the root tissue (Figure 4 and S5). However, blotting-based assays do not provide sufficient spatiotemporal resolution. For this reason, we constructed an *in situ* ROP sensor by inserting (i) ROP2 and (ii) the CRIB domain of the ROP effector RIC1 on opposite ends of a circularly permuted GFP (cpGFP). The cpGFP fluorescence should decrease when activated GTP-bound ROP2 interacts with the nearby CRIB domain (Figure S6D). Expression of CRIB-cpGFP-ROP2 and ROP2-mCherry from the same cassette yielded a ratiometric ROP sensor, which we called the CpGFP ROP Activity Probe (CRAP). As expected, the 561/488 nm CRAP ratio sensitively reported auxin (IAA, 10 nM) pulses in *CRAP;WT* roots in a microfluidic root chip setup (Figure S6E).

Strikingly, within 5 minutes of gravistimulation, CRAP-expressing roots developed asymmetric signal distribution with significantly more ROP activation at the lower side of the root (Figure S6F and Figure 5A). A GDP-locked CRAP (CRIB-cpGFP-ROP2^T20N^) failed to show this asymmetry, confirming that CRAP indeed reports ROP activation rather than e.g. local fluctuations of the cpGFP root microenvironment (Figure S6F). This identifies a novel asymmetric rapid response to gravity-induced auxin flux redirection in roots, as confirmed by the lack of CRAP asymmetry after inhibition of auxin transport by NPA (Figure S6I).

**Figure 5.**
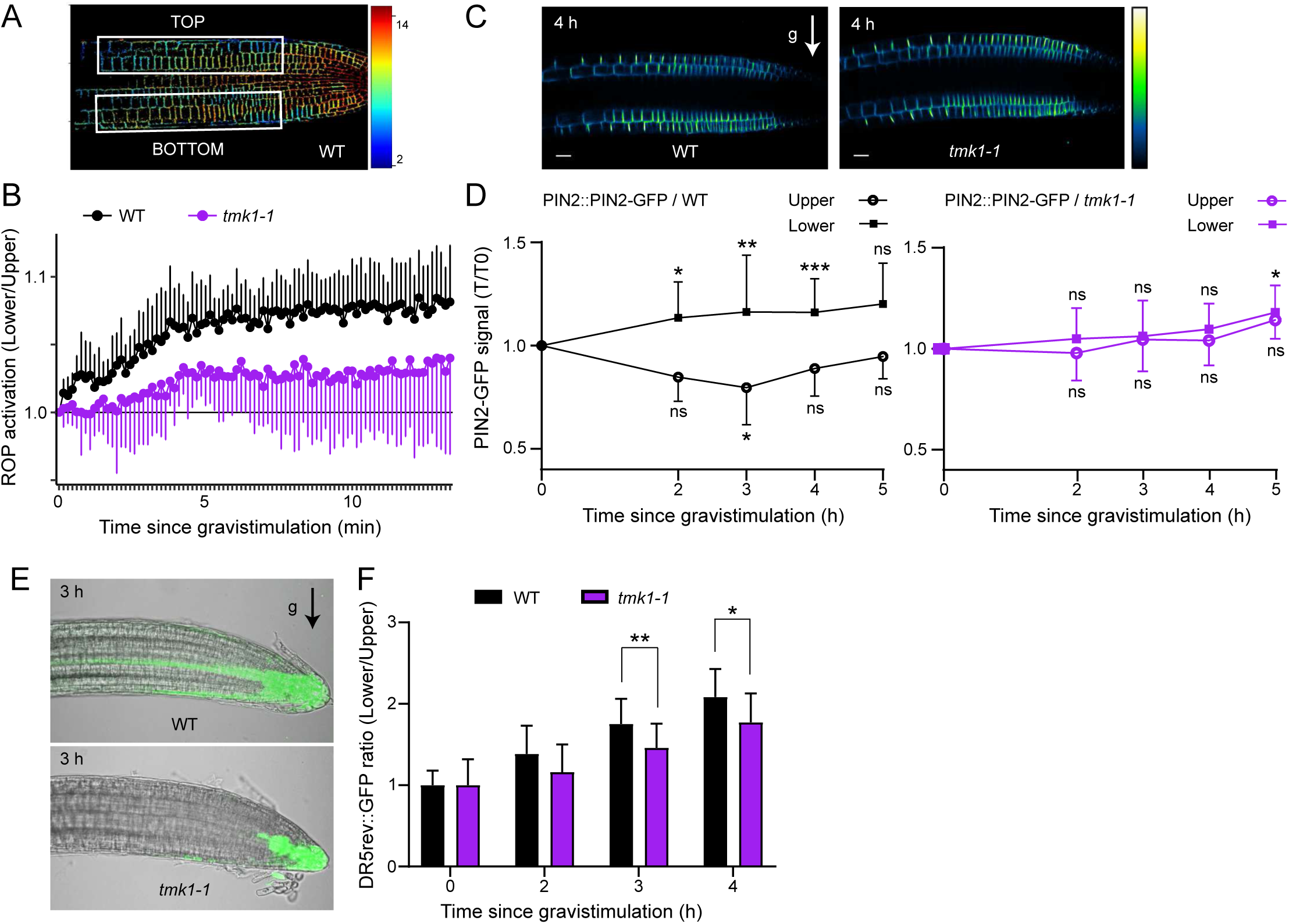
Asymmetric TMK1 activation for PIN2 asymmetry in root gravitropism. (A) Image of a root with strongly asymmetric ROP activity (reported by the CRAP sensor) in response to 15-minute gravistimulation. (B) Rapid gravistimulation-induced establishment of asymmetric ROP activity in WT or *tmk1-1* roots. (C) Representative images of *PIN2::PIN2-GFP* in 4-day-old WT and *tmk1-1* seedlings after 4 hours of gravistimulation. g, gravity vector. Scale bar, 20 μm. (D) Quantification of gravistimulation-induced PIN2-GFP asymmetry. Normalization to the first timepoint T0 for the upper or lower root side independently. Mean ± SD. Two-way ANOVA with Dunnett’s multiple comparisons, **** *p < 0.0001*. (E) Representative confocal images of asymmetric auxin response (visualized by *DR5rev::GFP*) at the lower side of the root after a 3-hour gravistimulation in WT and *tmk1-1* 4-day-old roots. Refer to Fig. S6H for corresponding images taken before gravistimulation. (F) Quantification of *DR5rev::GFP* asymmetry as a ratio of fluorescence intensity at the lower side to the upper side of gravistimulated roots at the indicated time points, normalized to the initial fluorescence value. Mean ± SD, n = 10. Two-way ANOVA with Dunnett’s multiple comparisons, \**p* < 0.05, \*\**p* < 0.01. g, gravity vector.

Given that auxin-induced ROP activation is TMK1-dependent (Figure 4 and S5), the gravitropic CRAP gradient likely mirrors asymmetric TMK1 activation by auxin flow from the root rip. Indeed, the *tmk1-1* mutation abolished asymmetric ROP activation in *CRAP;tmk1-1* roots (Figure 5B). These data collectively establish rapid asymmetric activation of the TMK1 kinase and downstream ROP signaling by redirection of auxin fluxes during gravitropic root bending.

### Asymmetric TMK1 activation mediates PIN2 asymmetry in root gravitropism

The rapid TMK1 activation along the lower root side corresponds well with the timeframe in which auxin induces PIN2 phosphorylation through TMK1 (Figure 1B), inferring that asymmetric TMK1 activation likely results in asymmetric PIN2 phosphorylation. Given that TMK1-regulated phospho-sites mediate PIN2 stability (see Figure 2), we decided to follow the fate of PIN2-GFP in gravistimulated roots. We observed asymmetric PIN2-GFP stabilization at the lower root side, as shown before^21,29,51^, and this was completely abolished in the *tmk1-1* mutant (Figure 5C and 5D).

We further assessed the role of TMK1 and its kinase activity by cloning a kinase-dead TMK1 construct carrying a mutation in the ATP-binding site and generating *UBQ10::TMK1^K616R^-mCherry* (TMK1^DN^) in a WT background. Notably, TMK1^DN^ expression perturbed the gravity-induced PIN2-GFP asymmetry (Figure S6G). Accordingly, it also conveyed an early defect in gravitropic root bending (Figure S6A and S6B), phenocopying the *tmk1-1* mutant (Figure 3A, S3F and S4D). The TMK1^DN^-expressing plants showed unperturbed levels of the endogenous TMK1 protein (Figure S6C), ruling out transgene-induced silencing of the endogenous *TMK1* gene. This shows that TMK1^DN^ causes a dominant negative phenotype, underscoring the importance of TMK1 kinase activity for both, gravity-induced PIN2 asymmetry and rapid bending response.

The requirement of TMK1 for the PIN2-GFP gradient suggests that TMK1 stabilizes PIN2 to enhance PIN2-mediated auxin flux from the root tip along the lower root side. To test this, we monitored the *DR5rev::GFP* auxin response reporter, which revealed a significantly decreased gravity-induced auxin asymmetry in *tmk1-1* compared to WT (Figure 5E and 5F, Figure S6H). Accordingly, inhibition of PIN-mediated auxin transport by NPA interfered not only with CRAP-reported asymmetric TMK1 activation (Figure S6I) but also the PIN2-GFP asymmetry (Figure S6J), confirming that polar auxin transport itself contributes to asymmetric TMK1 activation and subsequent PIN2 stabilization for further asymmetric auxin flow reinforcement. Finally, we followed PIN2-GFP gradient formation in the *abp1-TD1;abl3-1* background, which revealed a perturbed profile that was especially deficient in the maintenance phase (Figure S6K). The residual PIN2-GFP asymmetry contrasts with the complete abolishment in *tmk1-1* and likely indicates the existence of further unexplored ABL proteins upstream of TMK1.

Altogether, these data identify a positive feedback loop, in which, following gravistimulation, PIN2 redistributes auxin from the root tip to the lower root side, activating the TMK1 kinase, which promotes PIN2 phosphorylation and stabilization, channeling even more auxin along the lower root side and reinforcing the original gravity-induced auxin flow asymmetry.

### Auxin induces TMK1 interaction with PIN2 and phosphorylation of its hydrophilic loop

Our hitherto results demonstrate a strong functional relevance of TMK1-dependent PIN2 phosphorylation during root gravitropism. Co-localization of TMK1-GFP and PIN2-mCherry expressed from native promoters suggested a possibility for their direct interaction (Figure S7A). To test this, we immunoprecipitated TMK1-FLAG from *TMK1::TMK1-FLAG;tmk1-1* roots and used an anti-PIN2 antibody for detection of native PIN2. PIN2 did not co-immunoprecipitate with TMK1-FLAG in untreated samples. On the other hand, in auxin-treated roots (IAA, 5 or 20 nM, 15 or 30 minutes), PIN2 co-immunoprecipitated with TMK1-FLAG in a time-dependent and auxin concentration-dependent manner (Figure 6A and S7B). This suggests that auxin promotes the formation of a TMK1-PIN2 complex at the plasma membrane.

**Figure 6.**
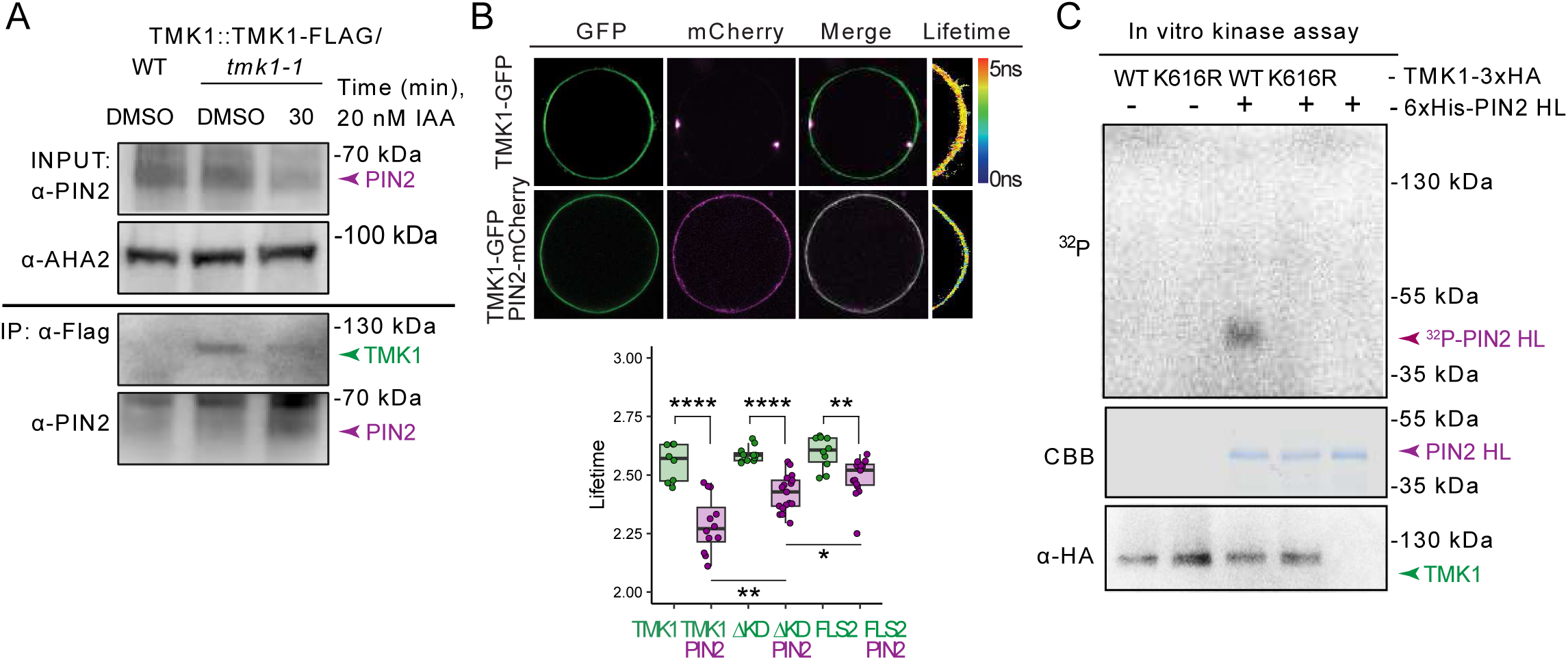
Auxin-mediated TMK1 interaction with and phosphorylation of PIN2. (A) Auxin promotes the interaction between TMK1 and PIN2 in a co-immunoprecipitation (Co-IP) assay. Microsomal protein fraction from 20 nM IAA- or DMSO-treated roots expressing *TMK1::TMK1-FLAG/tmk1-1* was immunoprecipitated with an anti-FLAG antibody, and endogenous PIN2 was detected by blotting with an anti-PIN2 antibody. See Figure S3A for Co-IP with 5 nM IAA where input PIN2 abundance in the protein microsomal fraction is increased. (B) FRET-FLIM analysis on transiently expressed *35S::TMK1-GFP, 35S::TMK1^ΔKD^-GFP, 35S::FLS2-GFP* and *35S::PIN2-mCherry* in root protoplasts. Fluorescence lifetime values are displayed as a heat map (for additional images see Figure S3B). One-way ANOVA with Holm-corrected post hoc tests, *p < 0.05, ****p < 0.0001. (C) *In vitro* kinase assay showing that TMK1-3xHA directly phosphorylates the 6His-PIN2 hydrophilic loop (HL). Top panel, ^32^P autoradiography; middle panel, CBB staining; bottom panel, immunoblot with an anti-HA antibody.

To verify the biochemical evidence for TMK1-PIN2 interaction, we performed fluorescence lifetime imaging on FRET-pair-tagged proteins (FRET-FLIM), a technique that quantitatively reports protein interactions. We introduced *35S::TMK1-GFP* and *35S::PIN2-mCherry* in Arabidopsis root protoplasts and measured the fluorescence lifetime of the GFP signal. PIN2-mCherry strongly reduced the lifetime of TMK1-GFP, demonstrating an interaction between TMK1 and PIN2 (Figure 6B). Notably, a truncated TMK1^ΔKD^-GFP variant without a kinase domain caused a significant ~35 % drop in the interaction strength compared to TMK1-GFP. The interaction of an unrelated receptor-like kinase FLS2-GFP with PIN2-mCherry was ~60 % weaker than that of TMK1-GFP (Figure 6B and S7C). These results establish both the contribution of the TMK1 kinase domain and the specificity of the TMK1-PIN2 interaction.

An auxin-induced TMK1-PIN2 interaction provides a plausible mechanism for the TMK1-dependent PIN2 phosphorylation observed in phospho-proteomic data (Figure 1 and S1). To test this, we performed an *in vitro* phosphorylation assay with ^32^P-ATP as a phosphate donor. We incubated a purified N-terminally HIS-tagged PIN2 hydrophilic loop (HIS-PIN2-HL) with TMK1-3HA immunoprecipitated from a root protein extract. The results showed that intact TMK1-3HA but not the kinase-dead version TMK1^K616R^-3HA was able to phosphorylate HIS-PIN2-HL (Figure 6C). We did not observe any auxin effect in this kinase assay (Figure S7D), presumably due to saturation of TMK1-3HA activity by endogenous auxin during immunoprecipitation from TMK1-3HA roots.

Taken together, these data demonstrate that the receptor-like kinase TMK1 interacts, in an auxin-dependent manner, with the PIN2 auxin efflux carrier and phosphorylates its hydrophilic loop to stabilize it.

## DISCUSSION

### Co-option of ancient auxin phospho-response for auxin feedback on its transport in vascular plants

Previous work indicated that while the rapid ABP1-TMK1-mediated auxin phospho-response is relevant for some rapid cellular auxin effects, specifically cytoplasmic streaming and apoplast acidification^13,17,18,52^, *abp1* and *tmk* mutants also show severe phenotypes in the long-term establishment of auxin- and auxin transport-positive channels after wounding and from externally applied auxin source, leading to vasculature formation^17^. The underlying mechanism of this so-called auxin canalization is largely unclear but at its center lies feedback regulation between auxin signaling and directional auxin transport^3,4^.

Here, mining of a root ABP1-TMK1 phospho-proteome revealed PIN auxin transporters as major phospho-targets of the ABP1-TMK1 cell-surface auxin perception. It follows that PIN phosphorylation by ABP1-TMK1 likely modulates directional auxin transport to delineate auxin channels for subsequent vascular differentiation and eventually other processes involving feedback regulation of auxin transport. Consistently, a 15-year-old model predicted extracellular auxin perception as a key signaling input parameter for auxin canalization^20^. Although the auxin phospho-response evolved in unicellular algae^18^, we find that it began targeting PINs only after the divergence of Bryophyta from the green lineage, likely in the common ancestor of vascular plants. The ABP1-TMK1-mediated PIN phosphorylation thus represents a recent evolutionary novelty that arose through the co-option of ancient rapid auxin response, presumably to enable the formation and regeneration of vasculature.

### ABP1-TMK1-mediated phosphorylation encodes PIN2 stability

Focusing on the dominant phospho-target PIN2, we report five ABP1-TMK1-dependent phospho-sites, the majority of which are induced by auxin and remarkably conserved. Strikingly, neither of these overlaps with previously published polarity-regulating PIN2 phospho-sites^38^. Indeed, both PIN2 stability and the physiological function of PIN2 in root gravitropism require ABP1-TMK1-dependent phospho-sites, implying the existence of two PIN2 phospho-codes; one for stereotypical maintenance of PIN2 polarity via AGC3 kinases^38^, and the other for dynamic adjustments of PIN2 stability in response to auxin.

### ABL3: Root-expressed auxin receptor acting redundantly with ABP1 in root gravitropism

Overall lack of *tmk1*-like phenotypes in *abp1* mutants, despite the strong similarity of phospho-proteomic signatures between *abp1* and *tmk1*^17^, contributed to the historical controversy surrounding ABP1. Indeed, while we confirmed an early gravitropic phenotype in the *tmk1* mutant, *abp1* mutant alleles show normal root bending, as reported before^42^. Hidden genetic redundancy with distant but structurally conserved ABL proteins has been invoked to explain this discrepancy, however, the recently identified ABL1 and ABL2 show minimal expression in the root, and the *abp1;abl1;abl2* triple mutant shows normal gravitropism^15^.

We identified the first root-expressed ABL protein, ABL3, through genetic redundancy with ABP1, and as an auxin binder and TMK1 interactor. Notably, ABL3 is paralogous to both ABL1 and ABL2 because they all belong to the 32-member Arabidopsis GLP family, which is distantly related to ABP1 by the ancient cupin fold^43^. ABP1 and ABL3 likely form part of an auxin-sensing complex docking on TMK1 in the root, providing a plausible model for auxin perception upstream of PIN2 phosphorylation.

While the field so far only scratched the surface of the real diversity of potential cell surface auxin receptors, this paints a picture in which specialized expression patterns of ABL auxin receptors confer specific functions on the rather ubiquitously expressed TMKs. There are likely more root-expressed ABLs awaiting discovery because *tmk1* and *abp1;abl3* mutants show a weaker phenotype than both the dominant-negative *ABP1::ABP1-5;abp1-TD1* line^15^ and higher-order *tmk* mutants^53,54^.

### Model for TMK1-based auxin feedback on PIN2-mediated transport in root gravitropism

The PIN2 transporter evolved as a specific component of efficient root gravitropism in seed plants^22^ and it is well documented that its abundance during gravistimulation becomes asymmetric with more PIN2 found at the lower root side^21,29^. This reinforces an initial auxin gradient and contributes to the robustness of root bending^30^. Nonetheless, the molecular mechanism behind this regulation has remained unknown since its discovery almost 20 years ago. Here, the wealth of our data together argues for a model encompassing auxin feedback on its transport.

A change in the gravity vector is sensed in the root columella, which establishes an initial auxin flow redirection to the lower side of the root^23^. This initial auxin asymmetry activates the TMK1 kinase specifically at the lower root side. Activated TMK1 then interacts with PIN2 in the epidermis and phosphorylates its hydrophilic loop at several conserved stability-regulating phospho-sites, leading to PIN2 stabilization in these cells. The resulting PIN2 abundance gradient further enhances auxin transport along the lower root side to the elongation zone, where it activates intracellular TIR1/AFB auxin signaling for growth inhibition and downward root bending. This demonstrates the existence of a positive feedback loop representing the first direct molecular mechanism for auxin feedback on its transport.

The TMK-based auxin feedback regulation likely represents a more general mechanism acting in various developmental contexts with different PIN transporters, thereby mediating specialized aspects of adaptive and self-organized plant growth and development. This is supported by the identification of TMK1-mediated PIN1 phosphorylation in the context of PIN polarity and auxin canalization^55^. The TMK-PIN mechanism evolved recently in vascular plants through the co-option of an ancestral auxin phospho-response from unicellular algae^18^ and it likely diversified to guide the auxin-mediated development of morphologically complex plants.

### Limitations of the study

Our work establishes a phosphorylation-based feedforward loop between ABP1/ABL3-TMK1 cell-surface auxin signaling and PIN2-mediated polar auxin transport in root gravitropism. Importantly, the newly identified root-expressed ABL3 auxin receptor extends the concept of ABP1/ABL signaling to roots, mirroring hypocotyl- and pavement cell-centered functions of the dominantly shoot-expressed ABL1 and ABL2 receptors^15^. Nevertheless, several lines of evidence indicate the existence of further unknown ABL proteins in the root. First, the gravitropic PIN2-GFP asymmetry is fully absent in *tmk1* but partially persists in *abp1;abl3* mutants. Second, *tmk1* and *abp1;abl3* mutants show a weaker root growth and bending phenotype than both the dominant-negative *ABP1::ABP1-5;abp1-TD1* line^15^ and higher-order *tmk* mutants^53,54^. Future studies should undertake a systematic effort to define which of the 32 Arabidopsis GLP proteins act as ABL auxin receptors. Furthermore, it remains a persistent mystery in the auxin transport field, how PIN phosphorylations on specific residues downstream of different signaling pathways^32^ selectively affect PIN activity, polarity, or stability.

## ACKNOWLEDGEMENTS

We thank W. Gray for providing material; N. Gnyliukh and E. Cervenova for help with manuscript preparation; J. Schmid for help with cloning. We thank Dolf Weijers, Mark Roosjen, and Andre Kuhn for discussions and support with phospho-proteomic analyses. We thank the Bioimaging and Life Science facilities at ISTA for their excellent service and assistance. The research leading to these results has received funding from the European Research Council (ERC) under the European Union’s Horizon 2020 research and innovation program grant agreement No 742985 and Austrian Science Fund (FWF): I3630-775 B25 to J.F; National Natural Science Foundation of China (Grant 32130010, 31422008), start-up funds from FAFU to T.X., Y.J. was funded by ERC no. 3363360-APPL under FP/2007-2013. L.R. was supported by FP7-PEOPLE-2011-COFUND ISTFELLOW program (IC1023FELL01) and the European Molecular Biology Organization (EMBO) long-term postdoctoral fellowship (ALTF 985-2016). S.T. was supported by the National Natural Science Foundation of China (32321001).

## AUTHOR CONTRIBUTIONS

Conceptualization: J.F., L.R.; Methodology: L.R., L.F. I.K.; Investigation: L.R., L.F., M.Z., C.G., D.V., A.M., M.R., Y.Y., Z.G., I.V., J.H., M.C., S.T., L.H., L.L., M.M.M.-B., Z.Q., G.M.; Resources: L.R., L.F., M.M.M.-B., Y.J., T.X.; Writing: J.F., L.F., L.R.; Review & Editing: all authors; Visualization: L.R., L.F.; Supervision: J.F., L.F., L.R.; Funding Acquisition: J.F.

## DECLARATION OF INTERESTS

The authors declare no competing interests.

## FIGURE LEGENDS

**Figure S1. Identification of PINs as ABP1-TMK1 phospho-targets by gene ontology analysis**

(A) Treemap of GO terms enriched for the ABP1-TMK1 phospho-proteome under mock conditions. Box sizes scale with −log10(p-value) (top) or fold enrichment (bottom).

(B) Treemap of GO terms enriched for the ABP1-TMK1 phospho-proteome under auxin treatment (100 nM IAA, 2 min) conditions. Box sizes scale with −log10(p-value) (top) or fold enrichment (bottom).

(C) Overview of phospho-peptides pertaining to PIN phospho-sites from Fig. 1A.

(D) Significant PIN1 and PIN3 phospho-site auxin profiles (FDR ≤ 0.01, 100 nM IAA).

(E) Localization of ABP1-TMK1-dependent phospho-sites on a ConSurf conservation-colored AlphaFold2 structure of PIN1.

(F) Localization of ABP1-TMK1-dependent phospho-sites on a ConSurf conservation-colored AlphaFold2 structure of PIN3.

**Figure S2. Supplementary data on PIN2-GFP phospho-lines**

(A) Normalized root angles after 12-hour gravistimulation of PIN2-GFP phospho-lines on medium with sucrose (1 %).

(B) Representative median root sections showing PIN2-GFP phospho-line polarity. Scale bar, 20 µm.

(C) Quantification of GFP signal in two independent PIN2^5-DEAD^-GFP phospho-lines compared to a PIN2^WT^-GFP phospho-line.

(D) qPCR quantification of *PIN2-GFP* mRNA in phospho-lines corresponding to (C). No *PIN2-GFP* was amplified in a wild-type Columbia-0 sample.

**Figure S3. Expression of *TMK* genes and analysis of the *tmk1-1* mutant**

(A) Relative transcription levels of *TMK1, TMK2, TMK3,* and *TMK4* in Arabidopsis roots from 803 samples assayed with the Affymetrix Arabidopsis ATH1 Genome Array. Data was obtained using the Genevestigator databases (https://genevestigator.com). Mean + SD.

(B) Root tip expression pattern of *TMK1*, *TMK2*, *TMK3*, and *TMK4* promoters fused to the GUS transcriptional reporter. Scale bar, 100 μm.

(C) Relative expression levels of *TMK1, TMK2, TMK3* and *TMK4* in different root cell types obtained from high-resolution spatiotemporal microarray analysis of 5-6-day-old roots reported in^56^ (ePlant http://bar.utoronto.ca/eplant).

(D) Representative intensity-colored image of a 4-day-old *TMK1::TMK1-GFP;WT* root.

(E) *TMK1* qPCR in *tmk1-1*.

(F) Root gravitropism profiles on medium without sucrose. Mean + or – SD.

**Figure S4. Characterization of ABL3 properties**

(A) Comprehensive expression profiles of *ABL3* and *ABP1/ABL/TMK* genes obtained using^57^.

(B) Snapshot of *ABL3* expression pattern obtained using ePlant/TAIR.

(C) *ABL3::GUS* staining (2h). Scale bar, 100 µm.

(D) Mutant root gravitropism profiles on medium without sucrose. Mean + or – SD.

(E) Representative images of mutant roots gravistimulated for 1 hour. Scale bar, 100 µm.

(F) Mutant root gravitropism profiles on medium without sucrose. Mean + or – SD.

(G) Multiple sequence alignment of ABP1/ABL protein sequences surrounding the auxin pocket with metal-coordinating residues (orange). Purple coloring highlights conservation.

(H) CETSA assay on *35S::ABL3-6xHIS-3xFLAG* seedlings in the presence of 100 µM IAA.

(I) Co-immunoprecipitation from Arabidopsis root protoplasts of ABL3-HA with TMK1-mCherry but not with anti-mCherry beads alone.

**Figure S5. Supplementary data for ROP activation through TMK1**

(A) Quantification of auxin effect on TMK1 phosphorylation in *TMK1::TMK1-FLAG;tmk1-1* roots assayed through TMK1-FLAG immunoprecipitation and Phos-tag Biotin Probe analysis. Phosphorylation levels detected with a Phos-tag Biotin Probe were normalized to signal obtained from anti-FLAG detection, and subsequently to the mock value. Mean ± SD of 3 biological replicates. Unpaired two-tailed t-test, \*\**p* < 0.01.

(B) Quantification of auxin effect on ROP6 and TMK1 levels in the WT or *tmk1-1* root microsomal protein fractions. Band intensities were normalized to the loading control, and subsequently to the first timepoint T0. Mean ± SD of 3 biological replicates. Two-way ANOVA with Dunnett’s multiple comparisons, \*\*\*\**p < 0.0001*.

(C) RT-qPCR of *TMK1* expression after auxin treatment. Normalization was done to *PP2A* as a reference gene. The expression level at the start of the experiment (T0) was set to a relative value of 1.

(D) Auxin effect on ROP2 activation assayed by native ROP pulldown from roots of the indicated genotypes. Full unedited blot relating to Fig. 4C.

(E) Quantification of gravistimulation effect on TMK1 phosphorylation in *TMK1::TMK1-FLAG;tmk1-1* roots assayed through TMK1-FLAG immunoprecipitation and Phos-tag Biotin Probe analysis. Phosphorylation levels detected with a Phos-tag Biotin Probe were normalized to signal obtained from anti-FLAG detection, and subsequently to the first timepoint T0. Mean ± SD of 3 biological replicates. One-way ANOVA with Dunnett’s multiple comparisons to timepoint 0 (\*\*\*\**p < 0.0001*).

(F) Quantification of gravistimulation effect on ROP6 and TMK1 levels in the WT or *tmk1-1* root microsomal protein fractions. Band intensities were normalized to the loading control, and subsequently to the first timepoint T0. Mean ± SD of 3 biological replicates. Two-way ANOVA with Dunnett’s multiple comparisons, \*\*\*\**p < 0.0001*.

**Figure S6. Requirement of TMK1 kinase activity for early root gravitropism**

(A) Representative images of primary roots expressing the *UBQ10::TMK1^K616R^-mCherry* construct.

(B) Root gravitropism profiles on medium without sucrose. Mean + or – SD.

(C) Western blotting shows that native TMK1 protein expression is not silenced in *UBQ10::TMK1^K616R^-mCherry* roots.

(D) Schematic showing the function of the CRAP sensor.

(E) CRAP sensor reports IAA pulses (10 nM IAA, highlighted in magenta) in a microfluidic root chip device.

(F) Dominant-negative mutation in the CRAP sensor abolishes rapid gravistimulation-induced establishment of CRAP asymmetry in WT roots. Mean + or – SD.

(G) Dominant negative effect of *UBQ10::TMK1^K616R^-mCherry* on gravistimulation-induced PIN2-GFP asymmetry. Mean + or – SD.

(H) Representative confocal images of *DR5rev::GFP;WT* and *DR5rev::GFP;tmk1-1* roots before gravistimulation.

(I) 10 µM NPA treatment abolishes rapid gravistimulation-induced establishment of CRAP asymmetry in WT roots. Mean + or – SD.

(J) 10 µM NPA treatment abolishes gravistimulation-induced PIN2-GFP asymmetry. Mean + or – SD.

(K) PIN2-GFP asymmetry defect in *abp1-TD1;abl3-1* double mutant. Mean + or – SD.

**Figure S7. Supplementary data on TMK1-PIN2 interaction**

(A) Co-localization of TMK1-GFP and PIN2-mCherry.

(B) Co-IP assay showing the interaction of TMK1 with PIN2 upon low auxin treatment (5 nM IAA). Microsomal protein fraction from WT was used as a control for unspecific binding of endogenous PIN2 (shown in Figure 3A, same experiment).

(C) FRET-FLIM analysis on transiently expressed *35S::TMK1^ΔKD^-GFP, 35S::TMK1^ΔKD^-GFP/35S::PIN2-mCherry, 35S::FLS2-GFP* and *35S::FLS2-GFP/35S::PIN2-mCherry* in root protoplasts (the same experiment is shown in Figure 3B). GFP fluorescence lifetime was calculated as described in the Methods section. The heat map depicts fluorescence lifetime values (see Figure 3B for lifetime analysis).

(D) *In vitro* kinase assay showing phosphorylation of the 6His-PIN2 HL substrate by TMK1-3xHA in the presence or absence of auxin. Top panel, ^32^P-autoradiography; middle panel, CBB staining; bottom panel, immunoblot with an anti-HA antibody.

## METHODS

### Molecular cloning, plant material, and growth conditions

All mutant alleles were in the *Columbia-0* (*Col-0*) background except *abp1-TD1*, which was in *Columbia-4* (*Col-4*). The T-DNA insertion lines *eir1-4* (SALK_091142) and *tmk1-1* (SALK_016360) were previously reported^21,58^, as was *abp1-C1* and *abp1-TD1*^59^. The T-DNA insertion line SAIL_441_G04, which harbors four insertions, was obtained from NASC and crossed with Col-0. Insertion-specific primers (Supplementary Table 3) were then used to out-segregate the SAILSEQ_441_G04.2 insertion in *ABL3*/AT4G14630 from SAILSEQ_441_G04.0 (AT4G09510), SAILSEQ_441_G04.1 (AT4G02930) and SAILSEQ_441_G04.3 (AT5G09590), yielding *abl3-1*. To obtain *abp1-C1;abl13-1* and *abp1-TD1;abl3-1*, we crossed the respective *abp1* mutants with *abl3-1*. The T-DNA insertion line SALK_203089C was obtained from NASC, verified (Supplementary Table 3), and used as *abl3-2*. The *abp1-TD1;abl3-2* double mutant was obtained by genetic crosses.

*PIN2::PIN2-GFP*, *DR5rev::GFP*, *TMK1::TMK1-FLAG*;*tmk1-1*, *TMK1::GUS*, *TMK2::GUS*, *TMK3::GUS*, and *TMK4::GUS* were described before^17,58,60,61^. We generated *PIN2::PIN2-GFP*;*tmk1-1* and *DR5rev::GFP*;*tmk1-1* by genetic crosses of the respective markers with *tmk1-1*. We generated *PIN2::PIN2-GFP*;*abp1-TD1;abl3-1* by a genetic cross with *abp1-TD1;abl3-1*. Shuta Asai kindly provided the *ABL3::GUS* and *35S::ABL3-6xHIS-3xFLAG* lines^62^. *DEX::TMK1^K616R^-HA* line (unpublished) was kindly provided by William Gray. To generate *TMK1::TMK1-GFP* and *35S::TMK1^ΔKD^-GFP*, the *TMK1* full length or *TMK1^ΔKD^* (amino acid 1-587) genomic DNA without a stop codon were amplified from *Col-0* DNA through PCR with *TMK1-FL-B1-F* and *TMK1-FL-B2-R/TMK1-ΔKD-B2-R* primers (Supplemental Table 3), respectively. The resulting *TMK1* sequences were inserted into pDONR221 by a BP reaction. Next, for the *TMK1::TMK1-GFP* construct, *pDONR P4-P1R pTMK1*, *pDONR221 gTMK1,* and *pDONR P2R-P3 GFP*; for *35S::TMK1^ΔKD^-GFP*, *pDONR P4-P1R p35S*, *pDONR221 gTMK1^ΔKD^* and *pDONR P2R-P3 GFP* were recombined into pB7m34GW vector by a MultiSite Gateway LR reaction. To generate *UBQ10::TMK1^K616R^-2xmCherry*, *TMK1^K616R^noSTOP/pDONR221* was obtained by site-directed mutagenesis, amplifying *pDONR221 TMK1* with the *TMK1KD-K616F* and *TMK1KD-K616R* primer pair. *pDONR221-TMK1^K616R^* was then recombined by LR reaction with *pDONR P4-P1R pUBQ10*^63^, *pDONR P2R-P3 2xmCHERRY-4xMyc*^64^ and pH7m34GW^65^ to obtain *UBQ10::TMK1^K616R^-2xmCherry* in pH7m34GW.

To clone PIN2 phospho-lines, Gibson Assembly (NEBuilder® HiFi DNA Assembly Master Mix, E2621L) was used to insert EGFP into the PIN2 coding sequence between Thr405 and Arg406 (according to^35^) and to assemble this fragment with an attL sites-containing pDONR221 backbone (Supplemental Table 3), yielding *pDONR221 PIN2^WT^-EGFP*. For *5-MIMIC* and *5-DEAD* constructs, we used gene synthesis (Integrated DNA Technologies, IDT) to obtain 1533-bp blocks containing EGFP and its flanking PIN2 CDS sequences with Ser179, Ser183, Thr233, Ser393, Ser439 triplets mutated to either GAC (Asp, *5-MIMIC*) or GCC (Ala, *5-DEAD*) (Supplementary table 3). These were Gibson-Assembled with a fragment amplified from the *pDONR221 PIN2^WT^-EGFP* plasmid (Supplementary table 3), yielding *pDONR221 PIN2^5-MIMIC^-EGFP* and *pDONR221 PIN2^5-DEAD^-EGFP*. Finally, the pDONR221 plasmids were recombined with pDONR-P4-P1R-pPIN2^22^ into pB7m24GW.3 by a multisite LR reaction (Gateway), and *eir1-4* (*pin2* null)^21^ was used for dipping.

The CRAP sensor was cloned using a combination of Gibson Assembly and the GreenGate approach. First, each GreenGate block was generated by fusing two PCR fragments – vector backbone and the respective CDS. ROP2 promoter fragment was subcloned into pGGA backbone fragment digested by BsaI. To obtain a non-ratiometric CRAP sensor, the following fragments were fused: pGGA-ROP2p. + pGGB-CRIB + pGGC-cpGFP + pGGD-ROP2 + pGGE009(UBI10term.) + pGGF-HYG. The resulting destination vector was sequenced and used as a template for cloning the entire CRAP CDS into a pGGD vector. This vector was then used in the GreenGate reaction to add the mCHERRY ratiometric control: pGGA-ROP2p + pGGB-mCHERRY-ROP2_UBI10term. + pGGC015(mCHERRY)-pGGD-CRAP-pGGE-HSP18.2term + pGGF005(HYG). ROP2-UBI10 fragment was amplified from the non-ratiometric CRAP sensor. Point mutagenesis to generate the dominant negative T20N mutation into ROP2 was performed as a single fragment Gibson Assembly with point-mutated compatible cohesive ends. The common building blocks were obtained from^66^ (pGGC015, pGGE-009, pGGF005). The pGGZ001 block with exchanged bacterial selection cassette (kanamycin) and pGGE-HSP18term was kindly provided by Dr. Andrea Bleckmann. The CRAP sensor was dipped into Col-0 to obtain *CRAP;WT* and this line was subsequently crossed with *tmk1-1* to obtain *CRAP;tmk1-1*.

To obtain seedlings co-expressing TMK1-GFP and PIN2-mCherry, we crossed *TMK1::TMK1-GFP;tmk1-1*^67^ with *PIN2::PIN2-mCherry;eir-1-4* (kind gift by Christian Luschnig) and imaged the F1 generation.

All constructs were transformed into the *Agrobacterium tumefaciens* strain pGV3101 by electroporation and further into plants by the floral dip method. The T2 generation was screened for single insertions and homozygous T3 lines were used for experiments.

Arabidopsis seeds were surface-sterilized with 70% (v/v) ethanol for 20 min, followed by commercial bleach (2.5% [v/v] sodium hypochlorite) containing 0.05% (v/v) Triton X-100 for 10 min, and finally washed four times with sterile water. Seed stratification was conducted in the dark at 4°C for 2 days. Unless indicated otherwise, seedlings were grown at 22°C on ½ MS plates with 1% agar and 1% sucrose, or in soil with 16h light/8h dark cycles, photoperiod at 80 to 100 mE m^−2^ sec^−1^.

### Bioinformatics

Phospho-proteomic analyses used data from^17,18,37^. Time-course profiles of auxin-induced phosphorylation were obtained with the AuxPhos tool (https://weijerslab.shinyapps.io/AuxPhos)^18^. Evolutionary rates for PIN1, PIN2, and PIN3 amino acids were calculated as ConSurf^68,69^ conservation scores and projected on the respective AlphaFold structures for visualization. Multiple sequence alignment of PIN2 orthologs (Uniprot IDs: Q9LU77, F6GXI9, E5KGD3, A0A251QTL1, A0A1D6P5D8, Q651V6, W1PK04, B5TXD0) or ABP1/ABLs (Uniprot IDs: P33487, P13689, P94040, P94072, Q9LEA7) was done with the MUSCLE tool available from EMBL-EBI^70^ and visualized in Jalview^71^.

For gene ontology enrichment, the “mock ABP1-TMK1 phospho-proteome” comprised Ensembl protein IDs of proteins concurrently hypo-phosphorylated in both *abp1-TD1* and *tmk1-1* mutants under mock conditions or—in the case of the “auxin-treated ABP1-TMK1 phospho-proteome”—auxin (2 min, 100 nM IAA) conditions^17^. These were submitted to the PANTHER extension of the TAIR database^72,73^ and queried for molecular function with all *Arabidopsis thaliana* proteins as a reference list. Annotation version and release date: [GO Ontology database DOI: 10.5281/zenodo.12173881 Released 2024-06-17]. Enrichment calculation was using Fisher’s Exact Test with a Bonferroni correction (p<0.05). The resultant terms were processed in REVIGO^74^ (parameters: medium list size of 0.7, clustering variables: p-value or fold enrichment, removal of obsolete GO terms, whole Uniprot reference set, SimRel semantic similarity) and visualized as treemaps with arbitrary coloring.

Plant pictograms were obtained from the Bioicons project under the MIT license. Attributions: Arabidopsis_thaliana icon by DBCLS https://togotv.dbcls.jp/en/pics.html is licensed under CC-BY 4.0 Unported https://creativecommons.org/licenses/by/4.0/. Zea_mays_cartoon icon by Daniel Carvalho https://figshare.com/authors/Plant_Illustrations/3773596 is licensed under CC-BY 4.0 Unported https://creativecommons.org/licenses/by/4.0/. DicotSeedling icon https://github.com/ginavong by Gina-Vong is licensed under CC0 https://creativecommons.org/publicdomain/zero/1.0/. The conifer branch pictogram was obtained from the free-for-use Pixabay repository.

The crystal structure of maize ABP1 (PDB ID: 1LRH) was superimposed with the AlphaFold2 structure of ABL3 (Uniprot ID: Q9LEA7) with the “super” command in Pymol. Tissue-specific expression profiles for *ABL1*, *ABL2*, *ABP1*, *ABL3*, *TMK1*, *TMK2*, *TMK3* and *TMK4* were compiled using a database of ~20,000 public Arabidopsis RNAseq experiments^57^.

### RT-qPCR

Total RNA was prepared from max. 100 mg of roots of 4-day-old seedlings according to the RNeasy Plant Mini Kit (Qiagen). cDNA was synthesized from 1µg of total mRNA using the QuantiNova Reverse Transcription Kit (Qiagen). Mutant expression analyses used 3-4 biological and 3 technical replicates pipetted into a 384-well plate using an automated JANUS Workstation (PerkinElmer). According to the manufacturer’s instructions, 5 µL reaction volume contained 2.5 µL Luna® Universal qPCR mastermix (NEB). RT-qPCR analyses were performed using the Real-time PCR Roche Lightcycler 480 and control PP2AA3 (At1G13320) primers were used as in^75^. For each of the evaluated genes, three different primer pairs were tested (Supplemental Table 3); except the *PIN2-GFP* phospho-line transgene, which was quantified using a single pair of primers without testing multiple sets.

### Fluorescence lifetime imaging

FRET-FLIM experiments were performed in protoplasts isolated from Arabidopsis root cell suspension as described previously^76^. 10 µg of plasmid DNA (*TMK1-GFP, TMK1^ΔKD-^GFP, PIN2-mCherry, FLS2-GFP*) were used for protoplast transfection, followed by incubation in a sterile 24-well microtiter plate overnight in the dark at room temperature. FRET-FLIM experiments were performed using a TriM Scope II inverted 2-photon microscope equipped with a FLIM X16 TCSPC Detector for time-correlated single photon counting (LaVision BioTec). Fluorescence lifetime image stacks (150 slices, with 0,082 ns time interval) were acquired. Image analyses were done in Fiji by performing a threshold mask from the sum projection of each stack and by averaging all the pixels at each time point of the stack. To yield an exponential decay with offset, the intensity at time point 0 was normalized.

### GUS staining

4-day-old seedlings were stained in 0.1 M sodium phosphate buffer (pH 7.0) containing 0.1% X-GlcA sodium salt, 1 mM K_3_[Fe(CN)_6_], 1 mM K_4_[Fe(CN)_6_] and 0.05% Triton X-100 for 20 min (*TMK1::GUS*), 2 h (*TMK4::GUS*, *TMK2::GUS*, *ABL3::GUS*) or 6 h (*TMK3::GUS*) at 37°C. Further, samples were incubated overnight in 80% (v/v) ethanol at room temperature. Tissue clearing was conducted as previously described^77^. DIC microscopy for analysis of the GUS staining assay was performed using an Olympus BX53 microscope equipped with 10x and 20x air objectives and a DP26 CCD camera.

### Root gravitropic assays

For sensitive phenotyping of the gravitropic response, unless indicated otherwise, seeds were germinated on sucrose-free ½ MS plates with 1% agar^26,78,79^. 5-day-old seedlings were transferred to fresh plates and incubated in a vertical position for 40 minutes for recovery. After rotation by 90°, the roots were imaged every 30 min on a vertical flatbed scanner (Epson Perfection V370 Photo). Image time series were stabilized using the StackReg Fiji plugin. Root curvature was analyzed with the Manual Tracking Fiji plugin and angles were calculated from root tip positions over time in Microsoft Excel.

### Imaging of transgenic lines

Confocal microscopy was performed on a vertical Zeiss LSM800 microscope^80^ equipped with a 20X Plan Apochromat air objective (NA = 0.8). GFP- and mCherry-tagged proteins [PIN2-GFP, TMK1-GFP, CRAP (mCherry/GFP), DR5rev::GFP] were excited at 488 and 561 nm, respectively, with emission collected in the following ranges: 490-576 nm or 560-700 nm, respectively.

PIN2-GFP phospho-lines were imaged by taking 12 Z-sections spanning the whole volume of each root. These were processed through “maximum intensity” projection in Fiji and total GFP signal was quantified across epidermal and cortical regions.

For imaging of *PIN2::PIN2-GFP* and *DR5rev::GFP* asymmetric distribution, 5-day-old seedlings were placed in a 1-well chambered coverglass (VWR, Kammerdeckglaser, Lab-Tek, Nunc, catalog number 734-2056) with a block of solid ½ MS medium^80^, optionally supplemented with mock or 10 µM NPA according to the experiment. For recovery, the chamber was incubated vertically in darkness for 2 h before imaging. Ten Z-sections spaced 1 µm for *PIN2::PIN2-GFP* and 4.5 µm for *DR5rev::GFP* were collected in the median root section before and after 90° rotation at the indicated time points. The “sum slices” intensity projection in Fiji was then applied. Marking epidermal and cortical regions together, the mean grey value was quantified as per^29^.

The CRAP sensor was validated by auxin treatments in a microfluidic vRootchip setup described previously^13,28^. For CRAP sensor imaging during gravistimulation, 5-day-old seedlings were placed vertically in a 1-well chambered coverglass with a block of solid 1/2 MS agar. The chamber was fitted inside a rotational stage. Seedlings were gravistimulated by turning the stage by 90°, achieving horizontal root position, and subsequently flipping the stage by 180°. Each root was imaged every 8.94 s for 15 min three times (with 180° flips in between) as technical replicates. GFP and mCherry signals were recorded simultaneously as a single track. Mean grey values at the lower and upper root sides were quantified, averaging the three technical replicates for each single root. Next, the 561/488 nm ratio indicative of ROP activity was calculated for the upper and lower root sides. Finally, to compare ROP activity between the two sides, the lower root side ratio was divided by the upper root side ratio and plotted over time. For visual representation, a root with a strong CRAP asymmetry was used. The last 10 time points of the 15-min imaging time course were averaged, and the pixel ratios reflecting CRAP activation were calculated as [mCherry - mCherry background] / [GFP - GFP background]. For plotting, the alpha value was derived from intensity and the Gaussian difference of the mCherry reference channel to suppress artifacts from numeric instability in low-intensity regions.

### Microsomal protein extraction

4-day-old-seedling roots were harvested, ground to powder in liquid nitrogen, and vortexed vigorously in extraction buffer 1 [50 mM Tris–HCl pH 7.5, 150 mM NaCl, Complete EDTA-free Protease Inhibitor cocktail (Roche), PhosSTOP phosphatase inhibitor cocktail (Roche)] in 1/10 (w/v) ratio. The resulting homogenate was centrifuged at 20000 g for 30 min at 4°C. The pellet was resuspended in extraction buffer 2 [50 mM Tris–HCl pH 7.5, 150 mM NaCl, 0.5% Nonidet P-40, 1% TritonX100, Complete EDTA-free Protease Inhibitor cocktail (Roche), PhosSTOP phosphatase inhibitor cocktail (Roche)], and centrifuged at 12000 g for 20 min at 4°C. Bradford assay was used to quantify protein in the supernatant. Samples were separated by SDS-PAGE (4-15% Mini-PROTEAN®TGX™ Precast Protein Gels (Bio-RAD)), transferred to a PVDF (polyvinylidene difluoride) membrane, and immuno-blotted using the following primary antibodies: affinity-purified TMK1 (1:1000)^58^, AHA2 (1:1000)^81^, PIN2 (1:1000)^21^ antibodies and anti-ROP6 (1:1000, Abiocode). The secondary antibody was anti-Rabbit-HRP (1:5000, GE Healthcare). Detection was done using the SuperSignal West Femto Maximum Sensitivity Substrate detection kit (ThermoFisher Scientific) and the Amersham 600RGB imager (GE Healthcare).

### CETSA

For CETSA from protoplasts, 10 µg of plasmid DNA (*35S::gGLP9-HA*) was used for protoplast transfection, followed by incubation in a sterile 24-well microtiter plate overnight in the dark at room temperature. The incubation buffer was exchanged for protein extraction buffer (50 mM Tris-HCl, pH 7.6, 150 mM NaCl, 10 % glycerol, 5 mM DTT, 1 mM PMSF, 0.5 % NP-40, complete Roche protease inhibitors) and protoplasts were lysed by 10 vigorous ice-vortexing(5s)-ice cycles, 30 min rotation at 4°C, and again 10 vigorous ice-vortexing(5s)-ice cycles. The lysate was centrifuged twice at maximum speed for 10 minutes (4°C) in a table-top centrifuge. The final supernatant represented the protein extract, which was first sampled for Western blotting and then split into two halves supplemented with either 100 µM IAA or mock treatment. IAA- or mock-treated extracts were incubated for 1 h on ice with occasional mixing. Next, the extracts were aliquoted for a 3 min incubation at temperatures between 42 and 47°C in a PCR machine (Bio-Rad) and returned to ice immediately. Finally, the samples were spun down in a tabletop centrifuge (12,000 rpm, 6 min, 4°C) and the supernatants were carefully transferred to new tubes for Western blotting (Anti-HA-HRP as described above).

CETSA from 5-day-old *35S::ABL3-6xHIS-3xFLAG* Arabidopsis seedlings used the same buffer and protocol with the only difference being the protein extraction procedure and the use of a different antibody for Western blotting (anti-FLAG^®^M2-Peroxidase (HRP), Sigma). Protein was extracted by adding ice-cold buffer to liquid-nitrogen-ground tissue, followed by centrifugation and supernatant collection as described for protoplast proteins.

### Western blot analysis of phosphorylated proteins

To analyze the phosphorylation status of TMK1 *in planta*, after corresponding treatment, 4-day-old *tmk1-1* roots expressing *TMK1::TMK1-FLAG* were ground to powder in liquid nitrogen and homogenized in ice-cold sucrose buffer [20 mM Tris–HCl pH 8, 0.33 M sucrose, Complete EDTA-free Protease Inhibitor cocktail (Roche), PhosSTOP phosphatase inhibitor cocktail (Roche)], followed by a centrifugation step at 5000 g for 10 min at 4°C. To obtain the membrane protein fraction, the supernatant was centrifuged at 20000 g for 30 min at 4°C and the resulting pellet was solubilized with lysis buffer [20 mM Tris–HCl pH 8, 150 mM NaCl, 1% TritonX100, Complete EDTA-free Protease Inhibitor cocktail (Roche), PhosSTOP phosphatase inhibitor cocktail (Roche)] and centrifuged at 20000 g for 10 min at 4°C. The corresponding supernatant was used for immunoprecipitation assay with anti-FLAG microbeads according to the manufactureŕs instructions (μMACS Epitope Tag Protein Isolation Kit (MACS Miltenyi Biotec)). Samples were separated by SDS-PAGE (4-15% Mini-PROTEAN®TGX™ Precast Protein Gels (Bio-RAD)) and transferred to a PVDF membrane, which was blotted with anti-FLAG^®^M2-Peroxidase (HRP) (Sigma) to detect total immunoprecipitated protein. Next, the membrane was stripped in 7 mL of mild stripping buffer (Abcam, 1 L: 15 g glycine, 1 g SDS, 10 mL Tween-20, pH 2.2) for 5 min, and the incubation was repeated with fresh buffer. The membrane was washed (2x 10 min in PBS, 2x 5 min in TBST) and re-blocked. The Phos-tag BTL-111 probe was then used according to the manufacturer’s instructions (www.wako-chem.co.jp) to detect phosphorylated protein levels.

### Co-immunoprecipitation assays

4-day-old *tmk1-1* seedlings expressing TMK1::TMK1-FLAG were treated with 5, 20 nM IAA or DMSO control for 30 minutes. Roots were harvested, ground to powder in liquid nitrogen, and subjected to protein microsomal fraction extraction. Solubilized proteins from microsomal fraction were immunoprecipitated using super-paramagnetic µMACS beads coupled to monoclonal anti-FLAG antibody according to the manufactureŕs instructions (Miltenyi Biotec). WT Col-0 extract was used as a control of the unspecific binding of endogenous PIN2. Proteins immunoprecipitated with anti-FLAG antibody were separated by SDS-PAGE (4-15% Mini-PROTEAN®TGX™ Precast Protein Gels (Bio-RAD)), transferred to a PVDF membrane and analyzed by immunoblot with anti-FLAG®M2-Peroxidase (HRP) (Sigma) antibody to detect TMK1-FLAG and with anti-PIN2 antibody to detect co-immunoprecipitated endogenous PIN2.

Tobacco leaves were infiltrated or co-infiltrated with overnight LB suspensions of *Agrobacterium tumefaciens* GV3101 carrying the desired expression plasmids (*35S::ABL3-HA*, *35S::ABL3-mCherry*, *35S::TMK1-HA*, *35S::TMK1-mCherry*). Including an overnight dark incubation, the infiltrated plants were grown for 36 hours, and leaves were subsequently harvested in liquid nitrogen. Frozen leaves were ground on ice in ice-cold extraction buffer (50 mM Tris-HCl, pH 7.6, 150 mM NaCl, 10 % glycerol, 5 mM DTT, 1 mM PMSF, 0.5 % NP-40, complete Roche protease inhibitors). The extracts were centrifuged in a tabletop centrifuge at top speed for 15 min and the supernatant was harvested, followed by a repetition of the same. The protein extracts were incubated with ChromoTek RFP-Trap Magnetic Agarose (rtma-20, Proteintech) for 1 hour at 4 ⁰C (rotating), washed 5 times with the extraction buffer, and analyzed by SDS-PAGE. Blotting was with an Abcam anti-mCherry antibody (ab167453) and anti-HA-HRP (described above).

### Active ROP assay on non-overexpressing plants

5-day-old seedling roots were harvested and incubated in ½ MS liquid medium with 100 nM IAA or mock treatment for 10 minutes and frozen in liquid nitrogen. Total protein was extracted in extraction buffer (25 mM HEPES pH 7.4, 100 mM KCl, 10 mM MgCl_2_, 1 mM PMSF, 5 mM Na3VO4, 5 mM NaF, 1 mM TCEP, cOmplete Protease Inhibitor Cocktail, Roche) in 2 steps. For every 100 mg of ground tissue, 400 μl of extraction buffer was added, then incubated with another 400 μl of extraction buffer containing 5% Triton X-100 for one hour. Active ROP proteins were pulled down using His-MBP-CRIB (BL21 *E*. *coli* total cell extract) conjugated to HisPurTM Ni-NTA Magnetic Beads (Thermo Fisher) by incubation of 200 μl total protein extract with 25 μl (initial volume) of beads for 2 hours at 4°C. The beads were washed with Washing buffer (25 mM HEPES pH 7.4, 300 mM KCl, 10 mM MgCl2, 12.5 mM imidazole) 3 times and boiled at 95°C with 30 μl SDS loading dye (BioRad). Protein samples were separated by 12% SDS-PAGE and analyzed by immune-blotting with an anti-ROP2 antibody (1:10,000, Abiocode). Input samples were blotted for total ROP2 content.

### Recombinant protein expression and purification from *E. coli*

The 6xHis-PIN2 HL recombinant protein was expressed using the 6xHis-PIN2 HL vector^82^ in *E. coli* BL21 (DE3) strain upon induction by 0.5 mM IPTG (Isopropyl β-D-1-Thiogalactopyranoside) at 16°C for 12 h. Cells were harvested by centrifugation at 4000 g for 15 min and washed with water. They were then lysed by sonication in lysis buffer (50 mM Tris-HCl pH7.5, 150 mM NaCl, 1 mM EDTA, 1 mM DTT, and 1% Triton X-100). The lysed solution was centrifuged at 12000 g for 15 min. The supernatant was then purified using Ni-NTA His affinity agarose (Thermo Scientific) according to the manufacturer’s instructions. 6xHis-PIN2 HL protein was eluted from the beads in elution buffer (50 mM Tris-HCl pH7.4, 150 mM NaCl, and 250 mM imidazole). Eluted protein samples were checked by SDS-PAGE and Coomassie brilliant blue staining (Bio-Safe™ Coomassie Stain, Bio-Rad). The protein concentration was determined with the Bradford method (Quick Start™ Bradford Reagent, Bio-Rad).

### TMK1-HA immunoprecipitation

To immunoprecipitate HA-tagged TMK1 protein, roots of 7-day-old *pUBQ10::TMK1-3xHA* and *DEX::TMK1^K616R^-HA* (previously induced for 48 h with 30 µM dexamethasone) seedlings were frozen and ground in liquid nitrogen, and subsequently homogenized in protein extraction buffer [PEB: 50 mM Tris-HCl pH7.5, 200 mM NaCl, 1% Triton X-100, 10 mM MgCl_2_, 1 mM MnCl_2_, Complete (Roche) protease cocktail and PhosSTOP phosphatase inhibitor cocktail (Roche)), followed by a 20 min centrifugation step at 14000 g, 4°C. In a fresh Eppendorf tube, 50 µL anti-HA agarose beads were added (Anti-HA Affinity Matrix, SIGMA; pre-washed with 200 µL PEB) to the supernatant. After 4 h of rotating at 4°C, the samples were spun down again at 2000 g, 4°C, and the supernatant was discarded. The agarose beads were washed twice with 200 µL PEB, and finally re-suspended into 50 µL PEB for further work.

### *In vitro* kinase assay

The *in vitro* kinase assay with [γ-^32^P]-ATP was conducted as reported^82^ with minor modifications. Immunoprecipitated TMK1-HA and TMK1^K616R^-HA (5 µL) from 7-day-old Arabidopsis seedlings, together with the recombinant 6xHis-PIN2 HL (10 µL) from *E*. *coli*, were added to the kinase reaction buffer (50 mM Tris-HCl pH 7.5, 10 mM MgCl_2_, 5 mM NaCl, 2.5 mM cold ATP (adenosine 5’-triphosphate), and 1 mM DTT) in the presence of 5 μCi [γ-^32^P]-ATP (NEG502A001MC; Perkin-Elmer). Reactions were incubated at 25°C for 90 min and terminated by adding 10 µL 5×SDS loading buffer. 20 µL reaction samples were then separated with 10% SDS-PAGE gel, developed with a phosphor-plate overnight. Eventually, the phosphor plate was imaged with a Fujifilm FLA 3000 plus DAGE system.

### Statistical analysis

Data processing and visualization was done in R version 4.0.4.

### Accession Numbers

Gene sequence data from this article can be found in the Arabidopsis Genome Initiative databases under the following accession numbers: AT4G02980 (*ABP1*), AT1G66150 (*TMK1*), AT1G24650 (*TMK2*), AT2G01820 (*TMK3*), AT3G23750 (*TMK4)*, AT5G57090 (*PIN2*), AT1G20090 (*ROP2*), AT4G35020 for (*ROP6*), AT4G30190 (*AHA2*), AT5G46330 (*FLS2*), AT1G72610 (*ABL1*), AT5G20630 (*ABL2*), AT4G14630 (*ABL3*).

## Notes

### Competing Interest Statement

The authors have declared no competing interest.

### Summary of Updates

The updated version reflects the project's progress and new findings since the first bioRxiv submission.

## REFERENCES

1. Friml, J. (2021). Fourteen Stations of Auxin. Cold Spring Harb. Perspect. Biol. 14, a039859. 10.1101/cshperspect.a039859.

2. Han, H., Adamowski, M., Qi, L., Alotaibi, S.S., and Friml, J. (2021). PIN-mediated polar auxin transport regulations in plant tropic responses. New Phytol. 232. 10.1111/nph.17617.

3. Berleth, T., and Sachs, T. (2001). Plant morphogenesis: Long-distance coordination and local patterning. Curr. Opin. Plant Biol. 4, 57–62. 10.1016/S1369-5266(00)00136-9.

4. Hajný, J., Tan, S., and Friml, J. (2022). Auxin canalization: From speculative models toward molecular players. Curr. Opin. Plant Biol. 65, 102174. 10.1016/J.PBI.2022.102174.

5. Adamowski, M., and Friml, J. (2015). PIN-dependent auxin transport: Action, regulation, and evolution. Plant Cell 27, 20–32. 10.1105/tpc.114.134874.

6. Luschnig, C., and Vert, G. (2014). The dynamics of plant plasma membrane proteins: PINs and beyond. Dev. 141, 2924–2938. 10.1242/dev.103424.

7. Park, J., Lee, Y., Martinoia, E., and Geisler, M. (2017). Plant hormone transporters: what we know and what we would like to know. BMC Biol. 15, 1–15. 10.1186/S12915-017-0443-X.

8. Péret, B., Swarup, K., Ferguson, A., Seth, M., Yang, Y., Dhondt, S., James, N., Casimiro, I., Perry, P., Syed, A., et al. (2012). AUX/LAX Genes Encode a Family of Auxin Influx Transporters That Perform Distinct Functions during Arabidopsis Development. Plant Cell 24, 2874–2885. 10.1105/tpc.112.097766.

9. Zhang, J., Nodzyński, T., Pěnčík, A., Rolčík, J., and Friml, J. (2010). PIN phosphorylation is sufficient to mediate PIN polarity and direct auxin transport. Proc. Natl. Acad. Sci. U. S. A. 107, 918–922. 10.1073/pnas.0909460107.

10. Wisniewska, J., Xu, J., Seifartová, D., Brewer, P.B., Růžička, K., Blilou, L., Rouquié, D., Benková, E., Scheres, B., and Friml, J. (2006). Polar PIN localization directs auxin flow in plants. Science (80-.). 312, 883. 10.1126/science.1121356.

11. Prigge, M.J., Platre, M., Kadakia, N., Zhang, Y., Greenham, K., Szutu, W., Pandey, B.K., Bhosale, R.A., Bennett, M.J., Busch, W., et al. (2020). Genetic analysis of the Arabidopsis TIR1/ AFB auxin receptors reveals both overlapping and specialized functions. Elife 9, e54740.

12. Qi, L., Kwiatkowski, M., Chen, H., Hoermayer, L., Sinclair, S., Zou, M., Del Genio, C.I., Kubeš, M.F., Napier, R., Jaworski, K., et al. (2022). Adenylate cyclase activity of TIR1/AFB auxin receptors in plants. Nature 611, 133–138. 10.1038/s41586-022-05369-7.

13. Li, L., Verstraeten, I., Roosjen, M., Takahashi, K., Rodriguez, L., Merrin, J., Chen, J., Shabala, L., Smet, W., Ren, H., et al. (2021). Cell surface and intracellular auxin signalling for H+ fluxes in root growth. Nature 599, 273–277. 10.1038/s41586-021-04037-6.

14. Fiedler, L., and Friml, J. (2023). Rapid auxin signaling: Unknowns old and new. Curr. Opin. Plant Biol. 75, 102443. 10.1016/j.pbi.2023.102443.

15. Yu, Y., Tang, W., Lin, W., Li, W., Zhou, X., Li, Y., Chen, R., Zheng, R., Qin, G., Cao, W., et al. (2023). ABLs and TMKs are co-receptors for extracellular auxin. Cell 186, 5457–5471.e17. 10.1016/j.cell.2023.10.017.

16. Sheen, J. (2024). The new horizon of plant auxin signaling via cell-surface co-receptors. Cell Res. 34, 343–344. 10.1038/s41422-023-00921-0.

17. Friml, J., Gallei, M., Gelová, Z., Johnson, A., Mazur, E., Monzer, A., Rodriquez, L., Roosjen, M., Verstraeten, I., Živanovič, B.D., et al. (2022). ABP1–TMK auxin perception for global phosphorylation and auxin canalization. Nature 609, 575–581. 10.1038/s41586-022-05187-x.

18. Kuhn, A., Roosjen, M., Mutte, S., Dubey, S.M., Carrillo Carrasco, V.P., Boeren, S., Monzer, A., Koehorst, J., Kohchi, T., Nishihama, R., et al. (2024). RAF-like protein kinases mediate a deeply conserved, rapid auxin response. Cell 187, 130–148.e17. 10.1016/j.cell.2023.11.021.

19. Mazur, E., Kulik, I., Hajný, J., and Friml, J. (2020). Auxin canalization and vascular tissue formation by TIR1/AFB-mediated auxin signaling in Arabidopsis. New Phytol. 226, 1375–1383. 10.1111/NPH.16446.

20. Wabnik, K., Kleine-Vehn, J., Balla, J., Sauer, M., Naramoto, S., Reinö Hl, V., Merks, R.M., Govaerts, W., and Friml, J. (2010). Emergence of tissue polarization from synergy of intracellular and extracellular auxin signaling. Mol. Syst. Biol. 6, 447. 10.1038/msb.2010.103.

21. Abas, L., Benjamins, R., Malenica, N., Paciorek, T.T., Wiřniewska, J., Moulinier-Anzola, J.C., Sieberer, T., Friml, J., and Luschnig, C. (2006). Intracellular trafficking and proteolysis of the Arabidopsis auxin-efflux facilitator PIN2 are involved in root gravitropism. Nat. Cell Biol. 8, 249–256. 10.1038/ncb1369.

22. Zhang, Y., Xiao, G., Wang, X., Zhang, X., and Friml, J. (2019). Evolution of fast root gravitropism in seed plants. Nat. Commun. 10, 1–10. 10.1038/s41467-019-11471-8.

23. Morita, M.T. (2010). Directional Gravity Sensing in Gravitropism. Annu. Rev. Plant Biol. 61, 705–720. 10.1146/annurev.arplant.043008.092042.

24. Friml, J., Wisniewska, J., Benkova, E., Mendgen, K., and Palme, K. (2002). Lateral relocation of auxin efflux regulator PIN3 mediates tropism in Arabidopsis. Nature 415, 806–809. 10.1038/415806a.

25. Kleine-Vehn, J., Ding, Z., Jones, A.R., Tasaka, M., Morita, M.T., and Friml, J. (2010). Gravity-induced PIN transcytosis for polarization of auxin fluxes in gravity-sensing root cells. Proc. Natl. Acad. Sci. U. S. A. 107, 22344–22349. 10.1073/pnas.1013145107.

26. Luschnig, C., Gaxiola, R.A., Grisafi, P., and Fink, G.R. (1998). EIR1, a root-specific protein involved in auxin transport, is required for gravitropism in Arabidopsis thaliana. Genes Dev. 12, 2175–2187. 10.1101/gad.12.14.2175.

27. Swarup, R., Friml, J., Marchant, A., Ljung, K., Sandberg, G., Palme, K., and Bennett, M. (2001). Localization of the auxin permease AUX1 suggests two functionally distinct hormone transport pathways operate in the Arabidopsis root apex. Genes Dev. 15, 2648–2653. 10.1101/gad.210501.

28. Fendrych, M., Akhmanova, M., Merrin, J., Glanc, M., Hagihara, S., Takahashi, K., Uchida, N., Torii, K.U., and Friml, J. (2018). Rapid and reversible root growth inhibition by TIR1 auxin signalling. Nat. Plants 4, 453–459. 10.1038/s41477-018-0190-1.

29. Baster, P., Robert, S., Kleine-Vehn, J., Vanneste, S., Kania, U., Grunewald, W., De Rybel, B., Beeckman, T., and Friml, J. (2013). SCFTIR1/AFB-auxin signalling regulates PIN vacuolar trafficking and auxin fluxes during root gravitropism. EMBO J. 32, 260–274. 10.1038/emboj.2012.310.

30. Retzer, K., Akhmanova, M., Konstantinova, N., Malínská, K., Leitner, J., Petrášek, J., and Luschnig, C. (2019). Brassinosteroid signaling delimits root gravitropism via sorting of the Arabidopsis PIN2 auxin transporter. Nat. Commun. 10, 1–15. 10.1038/s41467-019-13543-1.

31. Dubey, S.M., Han, S., Stutzman, N., Prigge, M.J., Medvecká, E., Platre, M.P., Busch, W., Fendrych, M., and Estelle, M. (2023). The AFB1 auxin receptor controls the cytoplasmic auxin response pathway in Arabidopsis thaliana. Mol. Plant 16, 1120–1130. 10.1016/j.molp.2023.06.008.

32. Tan, S., Luschnig, C., and Friml, J. (2021). Pho-view of Auxin: Reversible Protein Phosphorylation in Auxin Biosynthesis, Transport and Signaling. Mol. Plant 14, 151–165. 10.1016/J.MOLP.2020.11.004.

33. Zourelidou, M., Absmanner, B., Weller, B., Barbosa, I.C.R., Willige, B.C., Fastner, A., Streit, V., Port, S.A., Colcombet, J., van Bentem, S. de la F., et al. (2014). Auxin efflux by PIN-FORMED proteins is activated by two different protein kinases, D6 PROTEIN KINASE and PINOID. Elife 2014. 10.7554/ELIFE.02860.

34. Jia, W., Li, B., Li, S., Liang, Y., Wu, X., Ma, M., Wang, J., Gao, J., Cai, Y., Zhang, Y., et al. (2016). Mitogen-Activated Protein Kinase Cascade MKK7-MPK6 Plays Important Roles in Plant Development and Regulates Shoot Branching by Phosphorylating PIN1 in Arabidopsis. PLOS Biol. 14, e1002550. 10.1371/journal.pbio.1002550.

35. Vega, A., Fredes, I., O’Brien, J., Shen, Z., Ötvös, K., Abualia, R., Benkova, E., Briggs, S.P., and Gutiérrez, R.A. (2021). Nitrate triggered phosphoproteome changes and a PIN2 phosphosite modulating root system architecture. EMBO Rep. 22, 1–19. 10.15252/embr.202051813.

36. Ötvös, K., Marconi, M., Vega, A., O’Brien, J., Johnson, A., Abualia, R., Antonielli, L., Montesinos, J.C., Zhang, Y., Tan, S., et al. (2021). Modulation of plant root growth by nitrogen source-defined regulation of polar auxin transport. EMBO J. 40, 1–21. 10.15252/embj.2020106862.

37. Woudenberg, S., Alvarez, M.D., Rienstra, J., Levitsky, V., Mironova, V., Scarpella, E., Kuhn, A., and Weijers, D. (2024). Analysis of auxin responses in the fern Ceratopteris richardii identifies the developmental phase as a major determinant for response properties. Development 151. 10.1242/dev.203026.

38. Dhonukshe, P., Huang, F., Galvan-Ampudia, C.S., Mähönen, A.P., Kleinevehn, J., Xu, J., Quint, A., Prasad, K., Friml, J., Scheres, B., et al. (2015). Plasma membrane-bound AGC3 kinases phosphorylate PIN auxin carriers at TPRXS(N/S) motifs to direct apical PIN recycling. Dev. 142, 2386–2387. 10.1242/dev.127415.

39. Tena, G. (2023). ABP1’s new partners. Nat. Plants 9, 1941. 10.1038/s41477-023-01603-w.

40. Kuhn, A., and Weijers, D. (2024). Distant cousins come to ABP1’s rescue. Sci. China Life Sci. 67, 219–220. 10.1007/s11427-023-2498-0.

41. Dai, N., Wang, W., Patterson, S.E., and Bleecker, A.B. (2013). The TMK Subfamily of Receptor-Like Kinases in Arabidopsis Display an Essential Role in Growth and a Reduced Sensitivity to Auxin. PLoS One 8, 60990. 10.1371/journal.pone.0060990.

42. Gelová, Z., Gallei, M., Pernisová, M., Brunoud, G., Zhang, X., Glanc, M., Li, L., Michalko, J., Pavlovičová, Z., Verstraeten, I., et al. (2021). Developmental roles of Auxin Binding Protein 1 in Arabidopsis thaliana. Plant Sci. 303, 110750. 10.1016/j.plantsci.2020.110750.

43. Dunwell, J.M., Purvis, A., and Khuri, S. (2004). Cupins: The most functionally diverse protein superfamily? Phytochemistry 65, 7–17. 10.1016/j.phytochem.2003.08.016.

44. Chang, C., Eric Schaller, G., Patterson, S.E., Kwok, S.F., and Meyerowitz, E.M. (1992). The TMKl Gene from Arabidopsis Codes for a Protein with Structural and Biochemical Characteristics of a Receptor Protein Kinase. Plant Cell 4, 1263–1271. 10.1105/tpc.4.10.1263.

45. Oh, M.H., Wang, X., Kota, U., Goshe, M.B., Clouse, S.D., and Huber, S.C. (2009). Tyrosine phosphorylation of the BRI1 receptor kinase emerges as a component of brassinosteroid signaling in Arabidopsis. Proc. Natl. Acad. Sci. U. S. A. 106, 658–663. 10.1073/PNAS.0810249106.

46. Wang, X., Goshe, M.B., Soderblom, E.J., Phinney, B.S., Kuchar, J.A., Li, J., Asami, T., Yoshida, S., Huber, S.C., and Clouse, S.D. (2005). Identification and Functional Analysis of in Vivo Phosphorylation Sites of the Arabidopsis BRASSINOSTEROID-INSENSITIVE1 Receptor Kinase. Plant Cell 17, 1685–1703. 10.1105/TPC.105.031393.

47. Xu, T., Wen, M., Nagawa, S., Fu, Y., Chen, J.G., Wu, M.J., Perrot-Rechenmann, C., Friml, J., Jones, A.M., and Yang, Z. (2010). Cell surface- and Rho GTPase-based auxin signaling controls cellular interdigitation in Arabidopsis. Cell 143, 99–110. 10.1016/j.cell.2010.09.003.

48. Xu, T., Dai, N., Chen, J., Nagawa, S., Cao, M., Li, H., Zhou, Z., Chen, X., De Rycke, R., Rakusová, H., et al. (2014). Cell surface ABP1-TMK auxin-sensing complex activates ROP GTPase signaling. Science (80-.). 343, 1025–1028. 10.1126/science.1245125.

49. Smokvarska, M., Jaillais, Y., and Martinie, A. (2021). Function of membrane domains in rho-of-plant signaling. 663–681. 10.1093/plphys/kiaa082.

50. Pan, X., Fang, L., Liu, J., Senay-Aras, B., Lin, W., Zheng, S., Zhang, T., Guo, J., Manor, U., Van Norman, J., et al. (2020). Auxin-induced signaling protein nanoclustering contributes to cell polarity formation. Nat. Commun. 11, 1–14. 10.1038/s41467-020-17602-w.

51. Kleine-Vehn, J., Leitner, J., Zwiewka, M., Sauer, M., Abas, L., Luschnig, C., and Friml, J. (2008). Differential degradation of PIN2 auxin efflux carrier by retromer-dependent vacuolar targeting. Proc. Natl. Acad. Sci. U. S. A. 105, 17812–17817. 10.1073/PNAS.0808073105.

52. Lin, W., Zhou, X., Tang, W., Takahashi, K., Pan, X., Dai, J., Ren, H., Zhu, X., Pan, S., Zheng, H., et al. (2021). TMK-based cell-surface auxin signalling activates cell-wall acidification. Nature 599, 278–282. 10.1038/s41586-021-03976-4.

53. Dai, N., Wang, W., Patterson, S.E., and Bleecker, A.B. (2013). The TMK Subfamily of Receptor-Like Kinases in Arabidopsis Display an Essential Role in Growth and a Reduced Sensitivity to Auxin. PLoS One 8, 60990. 10.1371/journal.pone.0060990.

54. Marquès-Bueno, M.M., Armengot, L., Noack, L.C., Bareille, J., Rodriguez, L., Platre, M.P., Bayle, V., Liu, M., Opdenacker, D., Vanneste, S., et al. (2021). Auxin-Regulated Reversible Inhibition of TMK1 Signaling by MAKR2 Modulates the Dynamics of Root Gravitropism. Curr. Biol. 31, 228–237. 10.1016/J.CUB.2020.10.011.

55. Wang, J., Chang, M., Huang, R., Jiri, F., Yu, Y., Wen, M., Yang, Z., and Xu, T. (2022). Self-regulation of PIN1-driven auxin transport by cell surface-based auxin signaling in Arabidopsis. BioRxiv. 10.1101/2022.11.30.518523.

56. Brady, S.M., Orlando, D.A., Lee, J.Y., Wang, J.Y., Koch, J., Dinneny, J.R., Mace, D., Ohler, U., and Benfey, P.N. (2007). A high-resolution root spatiotemporal map reveals dominant expression patterns. Science (80-.). 318, 801–806. 10.1126/SCIENCE.1146265.

57. Zhang, H., Zhang, F., Yu, Y., Feng, L., Jia, J., Liu, B., Li, B., Guo, H., and Zhai, J. (2020). A Comprehensive Online Database for Exploring ~20,000 Public Arabidopsis RNA-Seq Libraries. Mol. Plant 13, 1231–1233. 10.1016/j.molp.2020.08.001.

58. Cao, M., Chen, R., Li, P., Yu, Y., Zheng, R., Ge, D., Zheng, W., Wang, X., Gu, Y., Gelová, Z., et al. (2019). TMK1-mediated auxin signalling regulates differential growth of the apical hook. Nature 568, 240–243. 10.1038/s41586-019-1069-7.

59. Gao, Y., Zhang, Y., Zhang, D., Dai, X., Estelle, M., and Zhao, Y. (2015). Auxin binding protein 1 (ABP1) is not required for either auxin signaling or Arabidopsis development. Proc. Natl. Acad. Sci. U. S. A. 112, 2275–2280. 10.1073/PNAS.1500365112.

60. Friml, J.J., Vieten, A., Sauer, M., Weijers, D., Schwarz, H., Hamann, T., Offringa, R., Jurgens, G., Cao, M., Chen, R., et al. (2003). Efflux-dependent auxin gradients establish the apical-basal axis of Arabidopsis. Nature 426, 147–153. 10.1038/nature02085.

61. Xu, J., and Scheres, B. (2005). Dissection of arabidopsis ADP-ribosylation factor 1 function in epidermal cell polarity. Plant Cell 17, 525–536. 10.1105/tpc.104.028449.

62. Asai, S., Cevik, V., Jones, J.D.G., and Shirasu, K. (2023). Cell-specific RNA profiling reveals host genes expressed in Arabidopsis cells haustoriated by downy mildew. Plant Physiol. 193, 259–270. 10.1093/plphys/kiad326.

63. Jaillais, Y., Hothorn, M., Belkhadir, Y., Dabi, T., Nimchuk, Z.L., Meyerowitz, E.M., and Chory, J. (2011). Tyrosine phosphorylation controls brassinosteroid receptor activation by triggering membrane release of its kinase inhibitor. Genes Dev. 25, 232–237. 10.1101/gad.2001911.

64. Simon, M.L.A., Platre, M.P., Assil, S., van Wijk, R., Chen, W.Y., Chory, J., Dreux, M., Munnik, T., and Jaillais, Y. (2014). A multi-colour/multi-affinity marker set to visualize phosphoinositide dynamics in Arabidopsis. Plant J. 77, 322–337. 10.1111/tpj.12358.

65. Karimi, M., Depicker, A., and Hilson, P. (2007). Recombinational cloning with plant gateway vectors. Plant Physiol. 145, 1144–1154. 10.1104/pp.107.106989.

66. Lampropoulos, A., Sutikovic, Z., Wenzl, C., Maegele, I., Lohmann, J.U., and Forner, J. (2013). GreenGate - A novel, versatile, and efficient cloning system for plant transgenesis. PLoS One 8. 10.1371/journal.pone.0083043.

67. Huang, R., Zheng, R., He, J., Zhou, Z., Wang, J., Xiong, Y., and Xu, T. (2019). Noncanonical auxin signaling regulates cell division pattern during lateral root development. Proc. Natl. Acad. Sci. U. S. A. 116, 21285–21290. 10.1073/PNAS.1910916116/SUPPL_FILE/PNAS.1910916116.SAPP.PDF.

68. Ashkenazy, H., Abadi, S., Martz, E., Chay, O., Mayrose, I., Pupko, T., and Ben-Tal, N. (2016). ConSurf 2016: an improved methodology to estimate and visualize evolutionary conservation in macromolecules. Nucleic Acids Res. 44, W344–W350. 10.1093/NAR/GKW408.

69. Yariv, B., Yariv, E., Kessel, A., Masrati, G., Chorin, A. Ben, Martz, E., Mayrose, I., Pupko, T., and Ben-Tal, N. (2023). Using evolutionary data to make sense of macromolecules with a “face-lifted” ConSurf. Protein Sci. 32, 1–12. 10.1002/pro.4582.

70. Madeira, F., Pearce, M., Tivey, A.R.N., Basutkar, P., Lee, J., Edbali, O., Madhusoodanan, N., Kolesnikov, A., and Lopez, R. (2022). Search and sequence analysis tools services from EMBL-EBI in 2022. Nucleic Acids Res. 10.1093/NAR/GKAC240.

71. Waterhouse, A.M., Procter, J.B., Martin, D.M.A., Clamp, M., and Barton, G.J. (2009). Jalview Version 2—a multiple sequence alignment editor and analysis workbench. Bioinformatics 25, 1189–1191. 10.1093/BIOINFORMATICS/BTP033.

72. Reiser, L., Subramaniam, S., Zhang, P., and Berardini, T. (2022). Using the Arabidopsis Information Resource (TAIR) to Find Information About Arabidopsis Genes. Curr. Protoc. 2, 1–43. 10.1002/cpz1.574.

73. Mi, H., Muruganujan, A., Casagrande, J.T., and Thomas, P.D. (2013). Large-scale gene function analysis with the panther classification system. Nat. Protoc. 8, 1551–1566. 10.1038/nprot.2013.092.

74. Supek, F., Bošnjak, M., Škunca, N., and Šmuc, T. (2011). REVIGO Summarizes and Visualizes Long Lists of Gene Ontology Terms. PLoS One 6, e21800. 10.1371/journal.pone.0021800.

75. Czechowski, T., Stitt, M., Altmann, T., Udvardi, M.K., and Scheible, W.R. (2005). Genome-wide identification and testing of superior reference genes for transcript normalization in arabidopsis. Plant Physiol. 139, 5–17. 10.1104/pp.105.063743.

76. Grones, P., Chen, X., Simon, S., Kaufmann, W.A., De Rycke, R., Nodzyński, T., Zažímalová, E., and Friml, J. (2015). Auxin-binding pocket of ABP1 is crucial for its gain-of-function cellular and developmental roles. J. Exp. Bot. 66, 5055–5065. 10.1093/jxb/erv177.

77. Malamy, J.E., and Benfey, P.N. (1997). Organization and cell differentiation in lateral roots of Arabidopsis thaliana. Development 124, 33–44. 10.1242/dev.124.1.33.

78. Xu, W., Ding, G., Yokawa, K., Baluška, F., Li, Q.F., Liu, Y., Shi, W., Liang, J., and Zhang, J. (2013). An improved agar-plate method for studying root growth and response of Arabidopsis thaliana. Sci. Rep. 3, 1–7. 10.1038/srep01273.

79. Thomas, M., Soriano, A., O’Connor, C., Crabos, A., Nacry, P., Thompson, M., Hrabak, E., Divol, F., and Péret, B. (2023). Pin2 Mutant Agravitropic Root Phenotype Is Conditional and Nutrient-Sensitive. Plant Sci. 329. 10.1016/j.plantsci.2023.111606.

80. von Wangenheim, D., Hauschild, R., Fendrych, M., Barone, V., Benková, E., and Friml, J. (2017). Live tracking of moving samples in confocal microscopy for vertically grown roots. Elife 6. 10.7554/eLife.26792.

81. Hayashi, Y., Nakamura, S., Takemiya, A., Takahashi, Y., Shimazaki, K., and Kinoshita, T. (2010). Biochemical Characterization of In Vitro Phosphorylation and Dephosphorylation of the Plasma Membrane H+-ATPase. Plant Cell Physiol. 51, 1186–1196. 10.1093/pcp/pcq078.

82. Tan, S., Abas, M., Verstraeten, I., Glanc, M., Molnár, G., Hajný, J., Lasák, P., Petřík, I., Russinova, E., Petrášek, J., et al. (2020). Salicylic Acid Targets Protein Phosphatase 2A to Attenuate Growth in Plants. Curr. Biol. 30, 381–395. 10.1016/j.cub.2019.11.058.

