## Supplements for "ABP1/ABL3-TMK1 cell-surface auxin signaling directly targets PIN2-mediated auxin fluxes for root gravitropism"

A

B

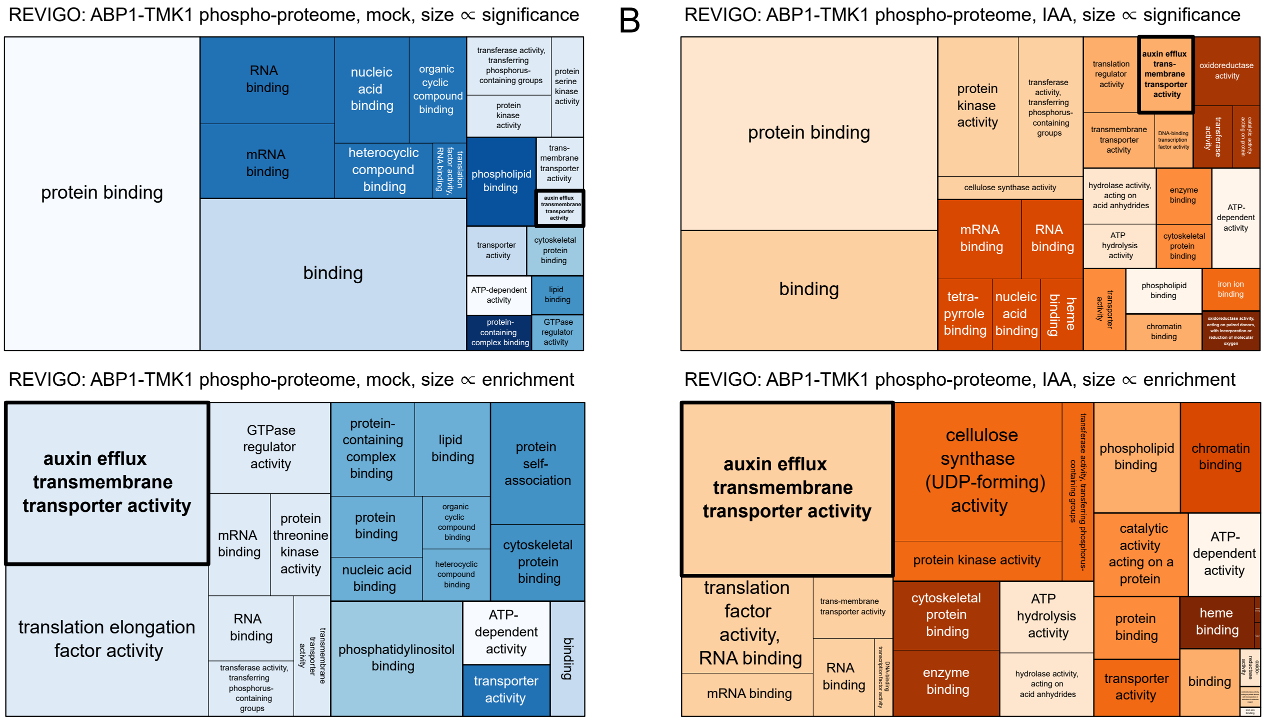


C

**PIN1[S271]** NpSNFGPGEAVFGSK

**PIN1[S337]** FHYQSGGSGGGGGA-HYPAPNPGMFpSPNT-GGGGGTAAK

**PIN2[S179]** VDpSDVISLNGR

**PIN2[S183]** VDSDVIpSLNGR

**PIN2[T233]** SHGGGLNSSMIpTPR

**PIN2[S393]** EMNMFVWSSSApSPV-SEANAK

**PIN2[S439]** AMQNLIENMpSPGRK

**PIN3[S366]** ELHMFVWSSNGpSPV-SDR

**PIN3[S389]** AGLNVFGGAPDNDQG-GRpSDQGAK

D

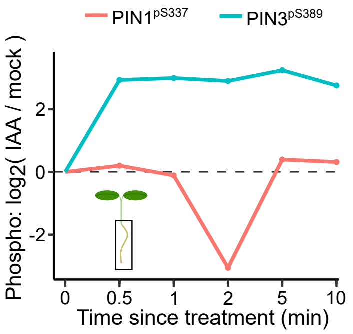


E

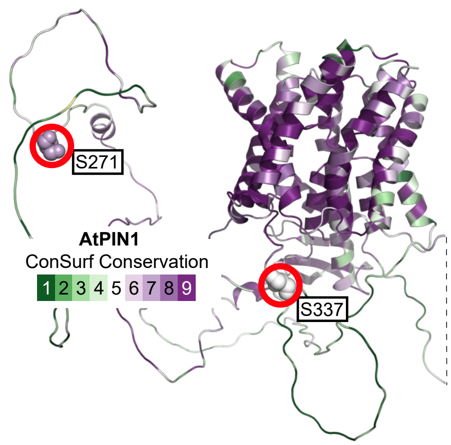


F

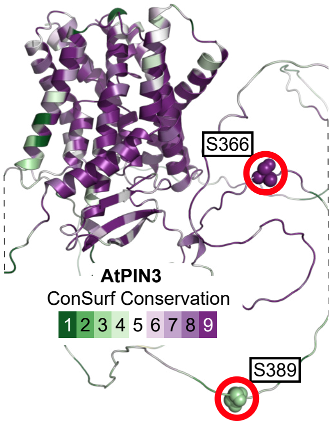


### Figure S1

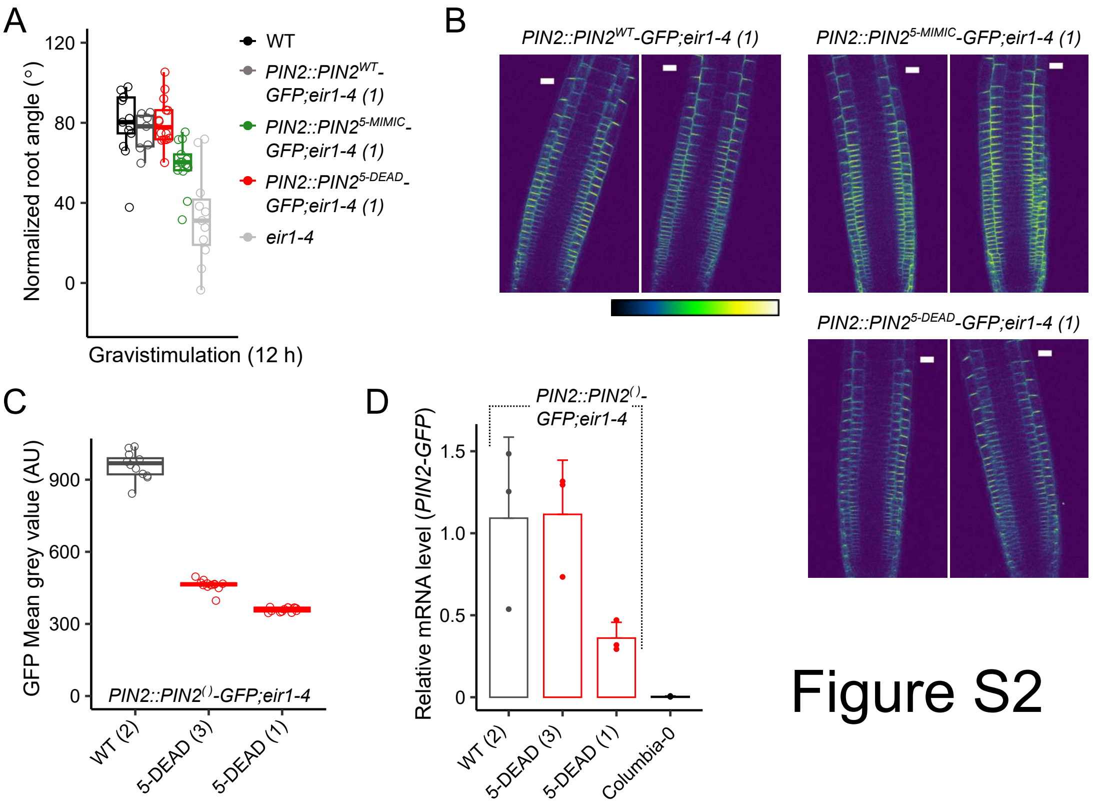
A
Normalized root angle (°)
120
80
40
0
Gravistimulation (12 h)
WT
PIN2::PIN2WT-GFP;eir1-4 (1)
PIN2::PIN25-MIMIC-GFP;eir1-4 (1)
PIN2::PIN25-DEAD-GFP;eir1-4 (1)
eir1-4
B
PIN2::PIN2WT-GFP;eir1-4 (1)
PIN2::PIN25-MIMIC-GFP;eir1-4 (1)
PIN2::PIN25-DEAD-GFP;eir1-4 (1)
C
GFP Mean grey value (AU)
900
600
300
0
PIN2::PIN2()-GFP;eir1-4
WT (2)
5-DEAD (3)
5-DEAD (1)
D
Relative mRNA level (PIN2-GFP)
1.5
1.0
0.5
0
PIN2::PIN2()-GFP;eir1-4
WT (2)
5-DEAD (3)
5-DEAD (1)
Columbia-0

### Figure S2

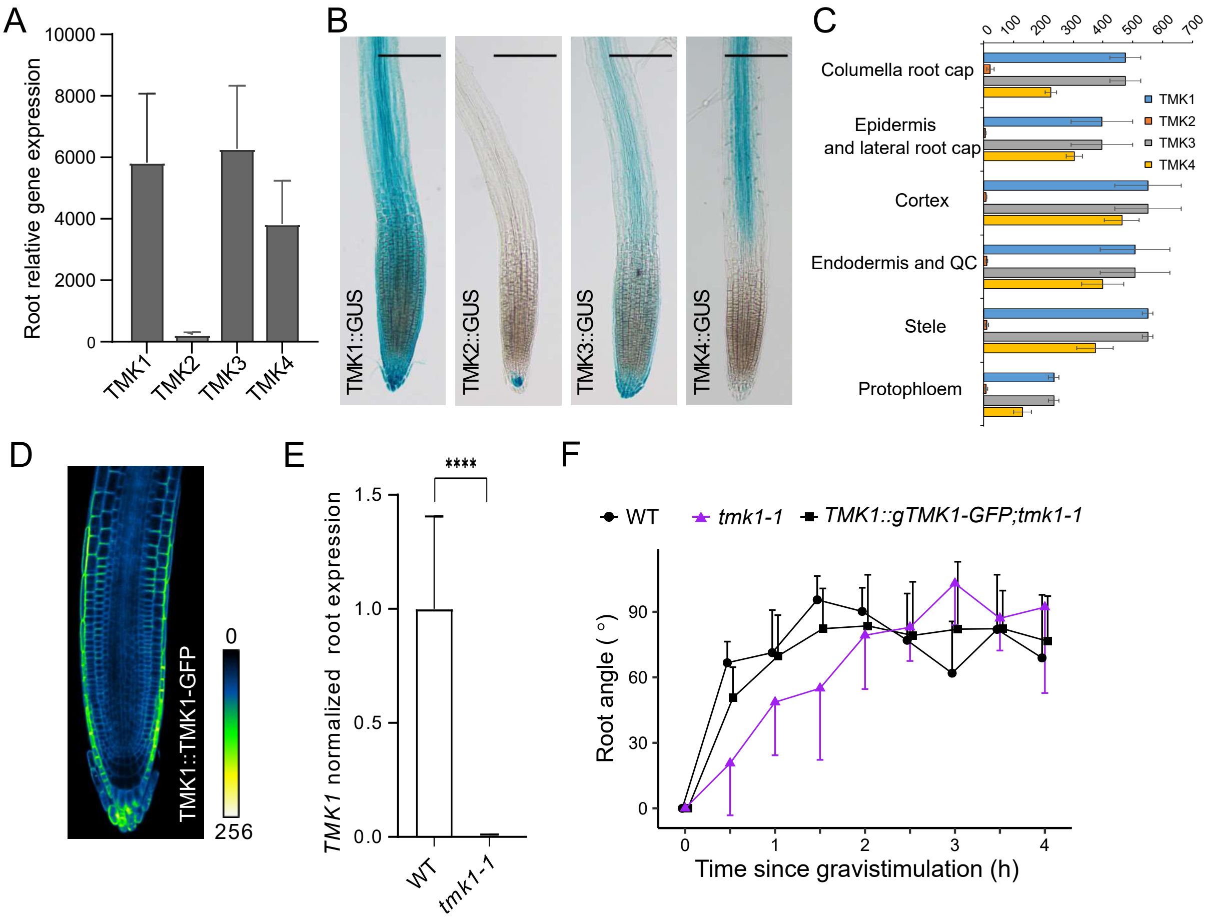

A
Root relative gene expression
10000
8000
6000
4000
2000
0
TMK1
TMK2
TMK3
TMK4
B
TMK1::GUS
TMK2::GUS
TMK3::GUS
TMK4::GUS
C
0
100
200
300
400
500
600
700
Columella root cap
Epidermis and lateral root cap
Cortex
Endodermis and QC
Stele
Protophloem
TMK1
TMK2
TMK3
TMK4
D
TMK1::TMK1-GFP
0
256
E
TMK1 normalized root expression
1.5
1.0
0.5
0.0
****
WT
tmk1-1
F
WT
tmk1-1
TMK1::gTMK1-GFP;tmk1-1
Root angle ( ° )
90
60
30
0
Time since gravistimulation (h)
1
2
3
4


### Figure S3

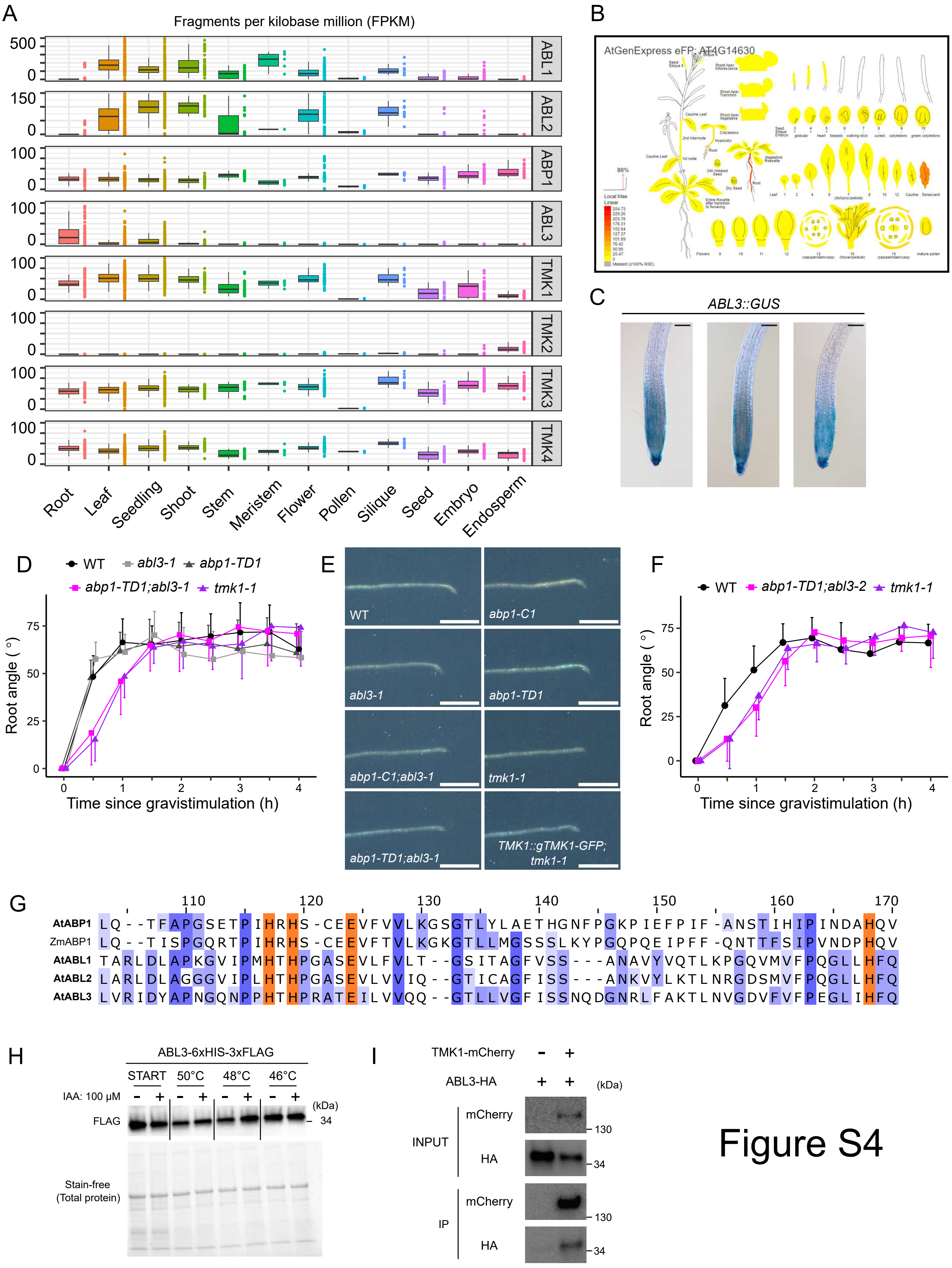

A
Fragments per kilobase million (FPKM)
ABL1
ABL2
ABP1
ABL3
TMK1
TMK2
TMK3
TMK4
Root
Leaf
Seedling
Shoot
Stem
Meristem
Flower
Pollen
Silique
Seed
Embryo
Endosperm
B
AtGenExpress eFP: AT4G14630
C
ABL3::GUS
D
WT
abl3-1
abp1-TD1
abp1-TD1;abl3-1
tmk1-1
Root angle ( ° )
Time since gravistimulation (h)
E
WT
abp1-C1
abl3-1
abp1-TD1
abp1-C1;abl3-1
tmk1-1
abp1-TD1;abl3-1
TMK1::gTMK1-GFP; tmk1-1
F
WT
abp1-TD1;abl3-2
tmk1-1
Root angle ( ° )
Time since gravistimulation (h)
G
AtABP1 LQ--TFAPGSETPIHRHS-CEEVFVVLKGSGTLYLAETHGNFPGKPIEFPIF-ANSTIHIPINDAHQV
ZmABP1 LQ--TISPGQRTPIHRHS-CEEVFTVLKGKGTLLMGSSSLKYPGQPQEIPFF-QNTTFSIPVNDPHQV
AtABL1 TARLDLAPKGVIPMHTHPGASEVLFVLT--GSITAGFVSS---ANAVYVQTLKPGQVMVFPQGLLHFQ
AtABL2 LARLDLAGGGVIPLHTHPGASEVLVVIQ--GTICAGFISS---ANKVYLKTLNRGDSMVFPQGLLHFQ
AtABL3 LVRIDYAPNGQNPPHTHPRATEILVVQQ--GTLLVGFISSNQDGNRLFAKTLNVGDVFVFPEGLIHFQ
H
ABL3-6xHIS-3xFLAG
START
50°C
48°C
46°C
IAA: 100 μM
(kDa)
FLAG
34
Stain-free (Total protein)
I
TMK1-mCherry
ABL3-HA
(kDa)
INPUT
mCherry
HA
IP
130
34


Figure S4

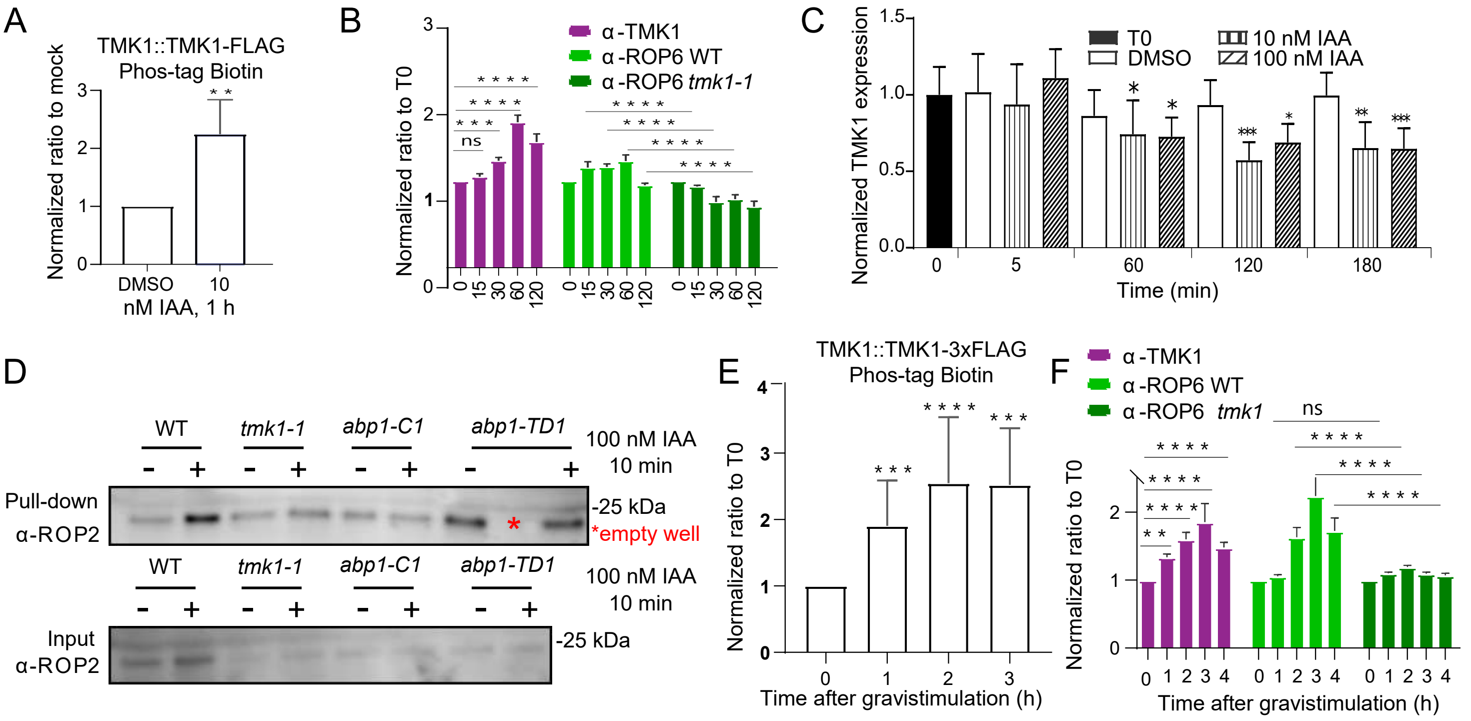
A
TMK1::TMK1-FLAG
Phos-tag Biotin
Normalized ratio to mock
DMSO
10
nM IAA, 1 h
B
Normalized ratio to T0
α -TMK1
α -ROP6 WT
α -ROP6 tmk1-1
C
Normalized TMK1 expression
T0
DMSO
10 nM IAA
100 nM IAA
Time (min)
D
WT
tmk1-1
abp1-C1
abp1-TD1
100 nM IAA
10 min
Pull-down
α-ROP2
-25 kDa
*empty well
Input
α-ROP2
E
TMK1::TMK1-3xFLAG
Phos-tag Biotin
Normalized ratio to T0
Time after gravistimulation (h)
F
α -TMK1
α -ROP6 WT
α -ROP6 tmk1
Normalized ratio to T0
Time after gravistimulation (h)

Figure S5

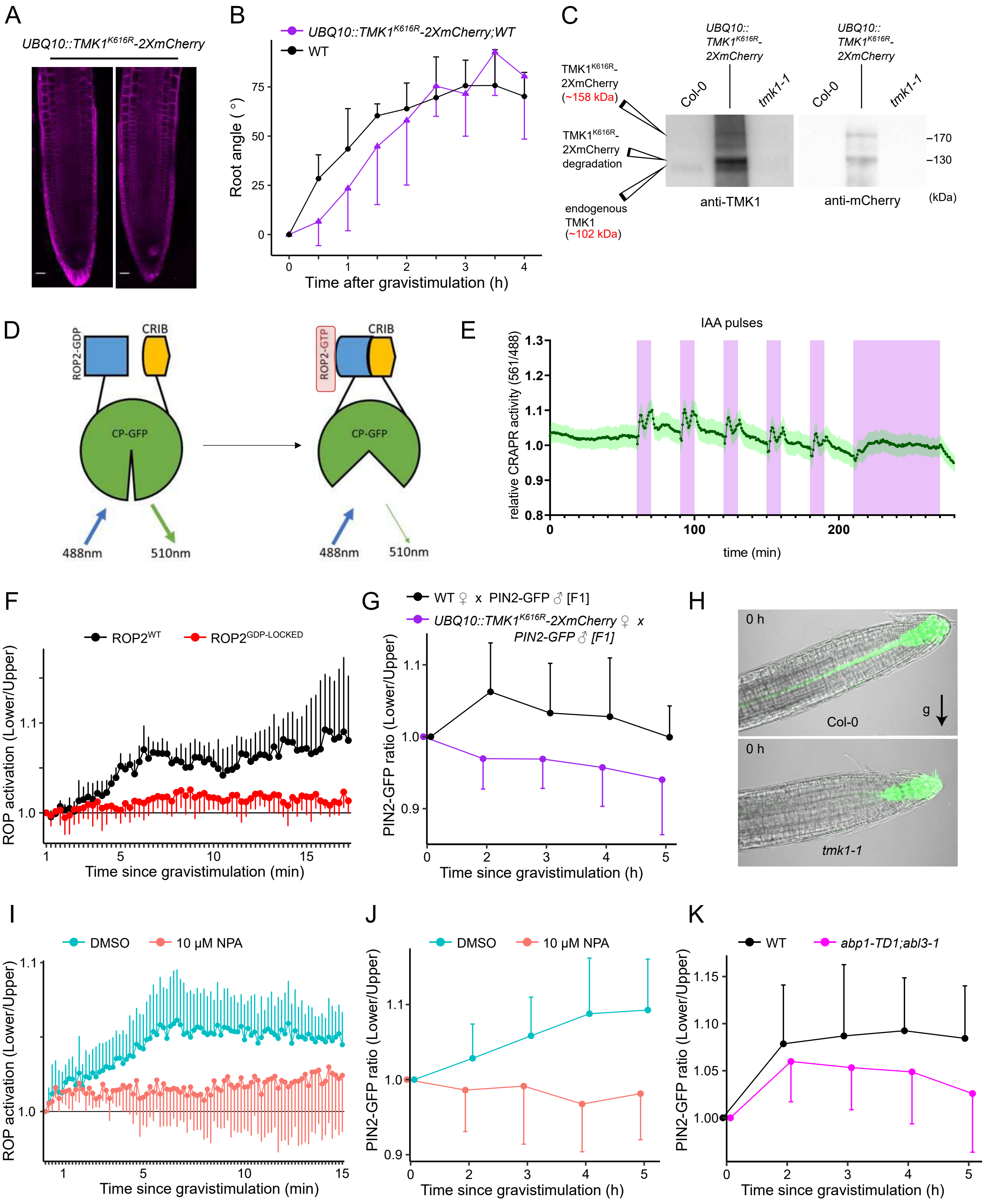
A
UBQ10::TMK1K616R-2XmCherry
B
UBQ10::TMK1K616R-2XmCherry;WT
WT
Root angle ( °)
Time after gravistimulation (h)
C
TMK1K616R-2XmCherry (~158 kDa)
TMK1K616R-2XmCherry degradation
endogenous TMK1 (~102 kDa)
UBQ10:: TMK1K616R-2XmCherry
Col-0
tmk1-1
anti-TMK1
anti-mCherry
(kDa)
D
ROP2-GDP
CRIB
ROP2-GTP
CP-GFP
488nm
510nm
E
IAA pulses
relative CRAPR activity (561/488)
time (min)
F
ROP2WT
ROP2GDP-LOCKED
ROP activation (Lower/Upper)
Time since gravistimulation (min)
G
WT ♀ x PIN2-GFP ♂ [F1]
UBQ10::TMK1K616R-2XmCherry ♀ x PIN2-GFP ♂ [F1]
PIN2-GFP ratio (Lower/Upper)
Time since gravistimulation (h)
H
0 h
Col-0
g
tmk1-1
I
DMSO
10 μM NPA
ROP activation (Lower/Upper)
Time since gravistimulation (min)
J
DMSO
10 μM NPA
PIN2-GFP ratio (Lower/Upper)
Time since gravistimulation (h)
K
WT
abp1-TD1;abl3-1
PIN2-GFP ratio (Lower/Upper)
Time since gravistimulation (h)

### Figure S6

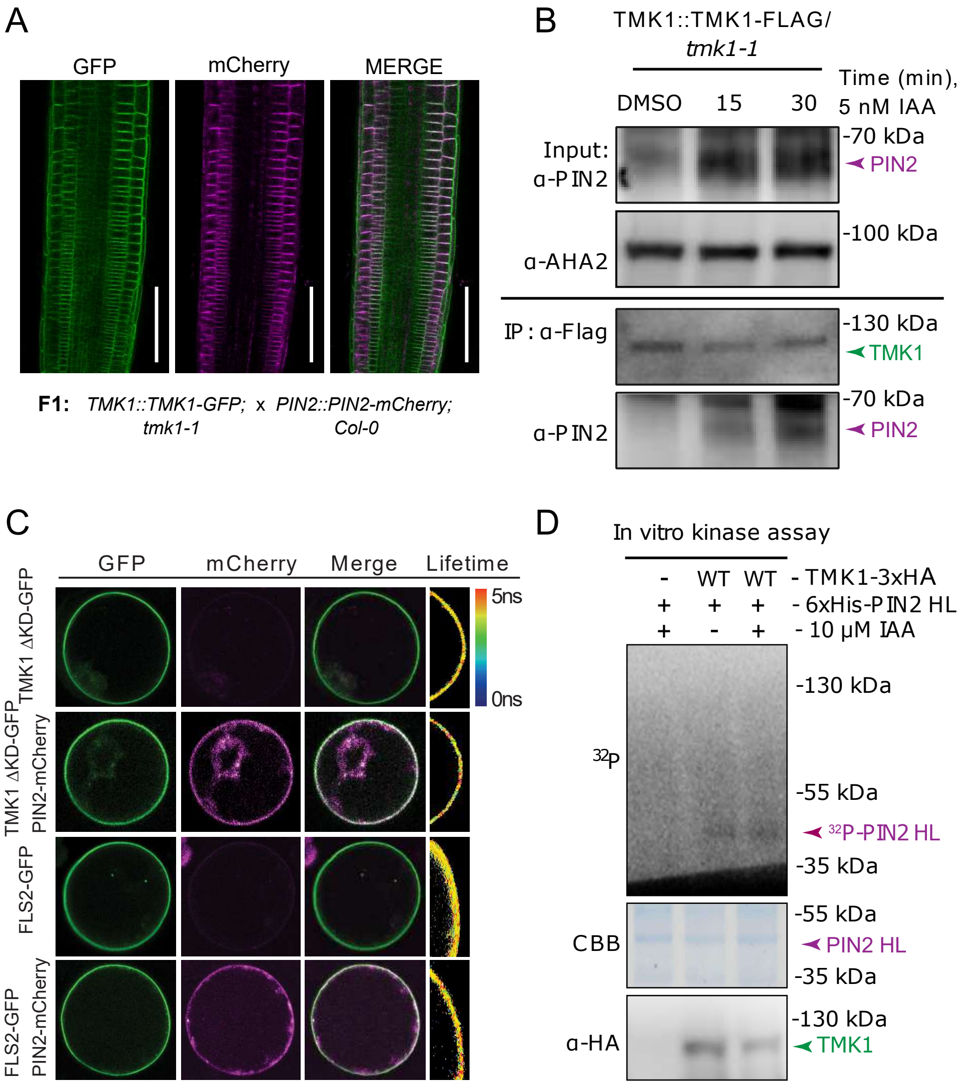
A
GFP
mCherry
MERGE
F1: TMK1::TMK1-GFP; tmk1-1 x PIN2::PIN2-mCherry; Col-0
B
TMK1::TMK1-FLAG/ tmk1-1
Time (min), 5 nM IAA
DMSO
15
30
Input: α-PIN2
-70 kDa
PIN2
α-AHA2
-100 kDa
IP : α-Flag
-130 kDa
TMK1
α-PIN2
-70 kDa
PIN2
C
GFP
mCherry
Merge
Lifetime
5ns
0ns
TMK1 ΔKD-GFP
TMK1 ΔKD-GFP PIN2-mCherry
FLS2-GFP
FLS2-GFP PIN2-mCherry
D
In vitro kinase assay
- WT WT - TMK1-3xHA
+ + + - 6xHis-PIN2 HL
+ - + - 10 μM IAA
32P
-130 kDa
-55 kDa
32P-PIN2 HL
-35 kDa
CBB
-55 kDa
PIN2 HL
-35 kDa
α-HA
-130 kDa
TMK1

Figure S7
